# Polarization MultiFocus Microscopy for volumetric super-resolution and orientation imaging of biofilaments

**DOI:** 10.1101/2025.11.19.687997

**Authors:** Louise Régnier, Caio Vaz Rimoli, Simli Dey, Feng-Ching Tsai, Guillermo A. Orsi, Sophie Brasselet, Bassam Hajj

**Affiliations:** Institut Curie, Université PSL, Sorbonne Université, CNRS UMR168, Physics of Cells and Cancer, Paris, France; Institut Curie, PSL Research University, INSERM U1339, CNRS UMR3666, Chemical Biology of Cancer Unit, Paris, France; Inria Center at University of Rennes, SAIRPICO Team, INSERM U1339, Institut Curie, Chemical Biology of Cancer Unit, Paris, France; Epigenetics and Chromatin Team, Institute for Advanced Biosciences, INSERM U1209, CNRS UMR5309, Université Grenoble Alpes, Grenoble, France; Aix Marseille Univ, CNRS, Centrale Med, Institut Fresnel, Marseille, France

## Abstract

Accessing molecular orientation in single molecule localization microscopy (SMLM) offers valuable insights into molecular ordering and organization in biological structures. Conventional single-molecule orientation-localization microscopy (SMOLM) methods typically rely on either engineering the microscope’s point-spread function (PSF) to encode the orientation information or on polarization resolved detection. While PSF engineering enables detailed orientation analysis, it often requires complex computational analysis and suffers from reduced performance in dense cellular environments due to PSF spreading and overlap. In contrast, polarization-based approaches are easier to implement and are more fit when imaging dense samples but are unable to retrieve the axial information of single molecules.

To overcome this limitation, we introduce the Polarization MultiFocus Microscope (PolMFM), a novel method for simultaneously retrieving the orientation and 3D position of single molecules. PolMFM combines the orientation measurement capabilities of a 4-polarization splitting scheme with a 3-planes multifocus microscope (MFM) enabling the reconstruction of molecular 2D orientation, wobble, and axial localization in a single acquisition. Through simulations, we demonstrate that PolMFM accurately recovers both orientation and 3D position, despite PSF defocusing. Experimental validation with reference samples shows that PolMFM matches the orientation precision of 4-Polar STORM, while uniquely adding axial information.

We demonstrate the power of PolMFM by resolving the orientation and 3D positions of molecules in actin filaments in fixed cells, and by revealing that chromatin in crickets undergoes major reorganization and increased ordering during spermiogenesis. These findings highlight the potential of PolMFM for high-precision, multidimensional super-resolution imaging in complex and crowded biological environments.

## Introduction

Single-molecule orientation-localization microscopy (SMOLM) has emerged as a powerful extension of single-molecule localization microscopy (SMLM), capable of resolving molecular ordering at the nanometer scale ^1–4^. In addition to localizing individual single molecules (SMs), as in conventional SMLM ^5–8^, SMOLM determines their orientation and rotational mobility. This added dimension reveals the role of molecular ordering in regulating biological function. By measuring the orientation of SMs, SMOLM can directly report on the orientation of a protein of interest (POI) when the fluorophore is rigidly bound^9,10^, or provide insights into the local environment’s composition and fluidity when the fluorophore’s motion is restricted—for example, in lipid bilayers ^11–14^. In both cases, the typical assumption of isotropic fluorophore emission and thus microscope’s symmetric point spread function (PSF) no longer holds. Instead, the limited mobility results in an asymmetric and polarization-dependent PSF pattern, due to the characteristic toroidal emission of a fixed dipole emitter ^15^.

SMOLM methods can be split into two main categories^16–18^. The first one uses the intrinsic or the engineered shape of the PSF to encode the position and orientation of SMs. PSF shape analysis, has first used defocused imaging ^19,20^, enabling the observation of the stepping of in vitro myosin on actin filaments ^21^. To enhance the encoding of orientation and wobbling information in the PSF, back focal plane (BFP) manipulation in phase ^13,22,23^ or polarization ^11,12,24^ has been proposed, with some examples using deep-learning for parameter estimation^25^ or phase mask design^26^. PSF engineering has shown unprecedented results in terms of precision and full 6D (3D localization, 3D orientation and wobbling) dipole parameters determination, especially when compact PSFs are achieved^11,12,22,24,26^. It has been successfully applied to study biological structures such as actin filaments^24^ or lipids in vitro and in cells^11^. However, this approach is sensitive to geometrical or polarization aberrations^12,22,24,26^, it also requires complex implementation^11^ and uses heavy computational analysis which requires fitting of features of the PSFs over multiple angular and position parameters.

The second category of SMOLM methods uses the polarized intensity dependence of the dipolar absorption or emission. Sequentially exciting the same SMs with multiple linearly polarized light states has enabled the retrieval of 2D-projected orientation in addition to spatial localization, - a strategy successfully demonstrated on in vitro DNA filaments^27^. Although this method remains simple to implement, it requires sequential acquisition of the same molecules making it impractical for most SMLM applications. Alternatively, the polarization state of the emission dipole can be accessed by projecting the emitted light onto distinct polarization directions, allowing simultaneous 2D (in-plane) orientation and localization measurements in a single acquisition.

Projecting the fluorescence emission onto two orthogonal polarization channels leads to orientation retrieval ambiguities^9^ that can be reduced by using a 4-polarization splitting scheme^10,28,29^. This configuration has enabled, for example, the tracking of 3D orientation and motion of quantum rod-conjugated myosin motor proteins (supposing a negligible wobble) ^28^, as well as the measurement of position, 2D orientation, and wobbling of actin filaments in various cellular actin networks^10^. A recent variant called POLCAM uses a polarization-resolved camera to acquire all four polarization channels in a single shot, offering a straightforward and compact implementation^30^.

In the 4-polarization detection scheme named 4Polar, the 2D localization of SMs relies on standard SMLM algorithms by combining orthogonal polarization channels. Orientation and wobbling are extracted through intensity-based analysis, without requiring complex PSF modeling or fitting procedures. Because the PSF footprint remains close to Gaussian shapes, this method is well-suited for long and densely labeled SMLM acquisitions. A recent extension of 4Polar to the retrieval of 3D orientation has been developed, combining 4-polarization projection with pupil plane reduction^31^.

While 4-polarization detection is interesting for its low level of experimental and computational complexity, a key limitation is its restriction to a single 2D imaging plane, since intensity measurement -based approaches are not compatible with depth encoding schemes usually performed in SMLM^32–34^. This precludes access to the axial distribution of molecular orientations, which limits the applicability of 4Polar SMOLM for three-dimensional volumetric biological imaging.

In this work, we extend the capability of 4Polar SMOLM to 3D spatial localization by combining 4-polarization detection^10^ with multifocus microscopy (MFM). We refer to this method as Polarization MultiFocus Microscopy (PolMFM). By a careful manipulation of the emitted wavefront as well as polarization projection, PolMFM enables the simultaneous retrieval of 2D-projected molecular orientation including wobbling, as well as 3D localization using only six images acquired simultaneously on a camera.

We demonstrate the measurement precision and capability of PolMFM on both 2D orientation and 3D localization estimations, using fluorescent molecules sparsely embedded in a polymer matrix, as well as on transiently bound Nile Red molecules to a lipid bilayer covering micrometric silica beads. We illustrate the applicability of PolMFM on the imaging of actin organization in fixed cells and on the spatial reorganization of the genome within the nucleus during spermiogenesis. These experiments highlight the system’s ability to provide axial information while preserving the performance of 4Polar SMOLM, extending its volumetric imaging range. Due to its relative simplicity, requiring only conventional Gaussian fitting for localization and intensity-based orientation analysis, PolMFM is readily applicable to a wide range of biological imaging scenarios. It provides a practical and powerful tool for generating high-precision, multidimensional super-resolution images in complex samples.

## Results

### Simultaneous 2D Orientation and 3D Localization of Single Molecules Using Polarized MultiFocus Microscopy

The goal in PolMFM is to access simultaneously the fluorophore’s orientation and wobbling projected in the sample plane, together with their 3D spatial localization, using SMs’ intensity measurements only. PolMFM combines 4-polarization splitting (4Polar) for determining the molecular 2D orientation and wobbling^10^, and multifocal microscopy (MFM) for the 3D localization^35^. Since MFM involves taking multiple images taken at different axial planes, we developed an optimized strategy that incorporates polarization projections among these different focal planes, shown in Fig. 1A.

**Figure 1:**
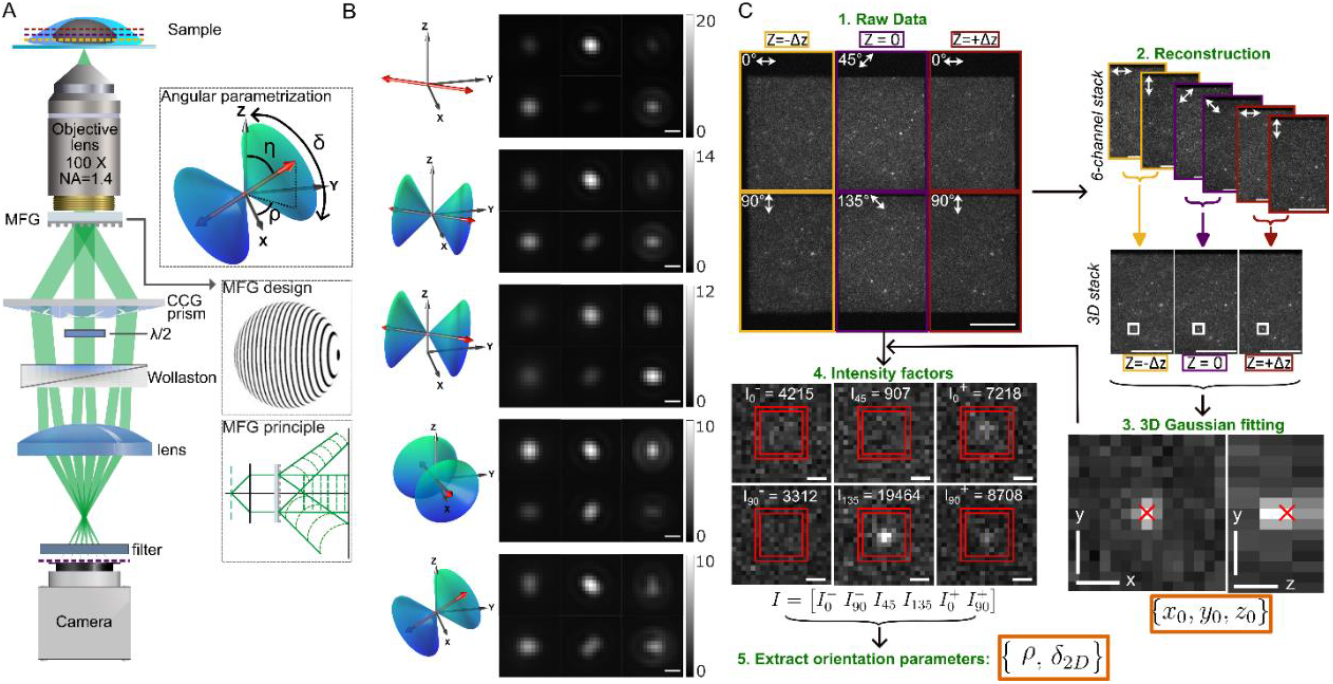
Single-molecule simulation and analysis in the PolMFM setup. A) Schematic of the polarization multifocus microscope. MFG: multifocus grating; CCG: chromatic correction grating. Insets: (top) angular parametrization of an electric dipole, modelling a fluorescent molecule; (middle) exaggerated view of the grating function of the 3-planes MFG; (bottom) schematics of the wavefronts at the different diffraction orders exiting the MFG. B) Simulated PSFs in the PolMFM, with different angular parameters, wobbling and axial position as depicted on the left. C) Overview of the SM analysis to retrieve localization and orientation parameters out of PolMFM raw data. (1) Raw data of JF549 dyes in PVA. Scale bar = 10 µm. (2) Reconstructed 6-channel stack (top). The 3D unpolarized z-stack is obtained by summing the perpendicular channels incoming from the same focal plane 2 by 2 (bottom). (3) A 3D gaussian fitting operation is performed on the 3D z-stack to infer the localization {*x*_0_, *y*_0_, *z*_0_}. Scale bar = 500 nm. (4) The intensity factors ***I*** are computing by summing the pixel intensities within a box (small red square) drawn around the SM position {*x*_0_, *y*_0_}, directly in the raw data. Prior to summation, the background, taken as the mean intensity value in the red ring around the box, is removed to the intensity values inside the box. (5) The orientation parameters {*ρ, δ*_2*D*_} are computed using equations (S19).

In PolMFM, the 3D spatial localization involves a 3-planes MFM scheme to generate an instantaneous 3D focal stack of the sample. These equally spaced focal planes are simultaneously imaged on a single camera by placing a custom-made diffraction grating and chromatic correction elements in the emission path of the microscope^35–37^(Fig. 1A). For SM 2D orientation determination, a Wollaston polarization beamsplitter projects each focal plane onto two perpendicular directions (0°, 90°). A half-wave plate is finally added in the optical path of the central plane, upstream of the Wollaston prism, to project this specific plane along the (45°, 135°) polarization directions. This results in four distinct polarization projections (0°, 90°, 45°, 135°) achieved across three focal planes (Fig. 1A). Given an axial spacing of around 400 nm between MFM planes (depending on the emission wavelength), each SM appears at least on two consecutive focal planes in PolMFM and is thus projected on 4 polarization channels.

To model PolMFM, fluorescent molecules are considered as electric dipoles 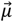 ^38^ whose orientation is described by a fast angular fluctuation within a cone of aperture *δ* and an averaged orientation (*ρ,η*) (Fig. 1A) ^39^. In this “rotation within a cone” model, the orientational behavior of SMs is encoded in the second-order dipole moment vector ***M*** ={*m*_*ij*_}, where *m*_*ij*_=**〈***µ*_*i*_*µ*_*j*_〉 and *i, j* ∈ {*x, y, z*}, **〈** 〉 being the average over the image integration time (Supplementary Note 1.2). Fig. 1B illustrates how SMs orientation, wobbling, and axial position affect the recorded images in PolMFM, in a simulation mimicking our experimental configuration. As wobbling increases (first two rows), the intensity differences between polarization channels decreases. Changes in axial position (third row) shifts the sharpest focal plane accordingly, indicating sensitivity to 3D spatial localization. In-plane orientation (fourth row) affects the balance of intensities across polarization channels, while out-of-plane orientation (last row) distorts the PSF asymmetrically. The retrieval of the 2D orientation and spatial parameters of SMs is based on the measurement of their 6-channels image vector ***I***, which we express as a matrix product between a 6×3 propagation matrix ***K***_**2D**_ and a reduced second-order dipole moment vector ***M***_**2D**_ that accounts only for the accessible moments (***M***_**2D**_ ={*m*_*xx*_, *m*_*yy*_, *m*_*xy*_}):

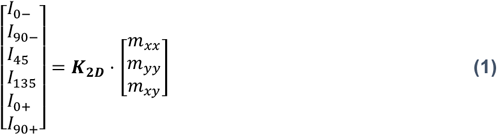

Here, ***K***_**2D**_ describes how the electric field vector propagates through the microscope (see Supplementary Note 1.4 for details).

To process the raw images, we implemented a multi-step analysis pipeline (Fig. 1C). In the first step, each raw image (Fig. 1C-1) is converted into a 6-channel stack (Fig. 1C-2) using a pre-calibration procedure based on a z-stack of immobilized 100 nm fluorescent beads. This calibration was used to align the channels and determine the interplane spacing (Supplementary Note 4.1).

Next, to estimate the spatial 3D position of each SM, unpolarized images are generated for each focal plane. For this, perpendicular-polarization (0°-90° or 45°-135°) channels corresponding to each focal plane are summed (Fig. 1C-2). Standard SMLM algorithms are then applied to detect single molecules in these reconstructed 3-focal plane volume and their 3D coordinates {*x*_0_, *y*_0_, *z*_0_} are determined using 3D Gaussian fitting^35^ (Fig. 1C-3). The lateral position {*x*_0_, *y*_0_} of each detected SM is used as the center of a 11×11 pixels-ROI, within which pixel intensities are summed to yield the 6-channels integrated intensity vector ***I*** =[*I*_0−_ *I*_90−_ *I*_45_ *I*_135_ *I*_0+_ *I*_90+_]^*T*^ (Fig. 1C-4). The background for each ROI is estimated by an averaging over the outer periphery pixels of the ROI and subtracted before summation. Note that intensity imbalance between polarization channels resulting from polarization aberrations in the setup, is corrected for using an unpolarized white lamp image (Supplementary Note 4.2).

Finally, the second-order moments *m*_*ij*_(*i, j* ∈ {*x, y*}) are retrieved by the pseudo-inversion operation 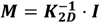 (Fig. 1C-5). The propagation matrix ***K***_**2D**_ used here is obtained experimentally, using a calibration procedure that mimics dipoles of various orientations (Supplementary Note 4.3). The in-plane orientation parameters {*ρ, δ*_2*D*_} are finally extracted from the second-order moments *m*_*ij*_ using the procedure developed in ^10^ (Supplementary Note 1.5).

### Simulations

A known challenge in polarized localization microscopy is that out-of-focus PSFs can deviate significantly from the ideal Airy pattern under restricted dipole wobbling, potentially affecting localization precision^40,41^. To evaluate whether PSF defocusing in the different channels compromises the accurate extraction of polarization-resolved intensities, we conducted simulations comparing PolMFM to conventional 4-polarization where all four polarization channels are recorded in a single focal plane. The simulation conditions are performed using a photon budget of 1000 photons distributed across the six PolMFM channels (or the four 4-polar channels respectively) and moderate noise levels (2 background photons per pixel) (Material and Methods). These simulation results, performed for various orientation parameters and defocus conditions, are shown in Fig. 2. Despite PSF defocusing, both PolMFM and the 4Polar configuration exhibited comparable performances on the orientation parameters retrieval (Fig.2A). The precision obtained on the in-plane orientation estimation remained below 5° for *δ* = 1°, increased to ∼13° at *δ* = 110°, and worsened as wobbling approached 180°, which is expected since in-plane orientation becomes ill-defined. When molecules were slightly tilted out of plane (*η* = 60°), errors increased moderately but equally in both systems (Fig. 2B). Similarly, biases and errors on the wobbling *δ*_2*D*_ retrieval exhibit similar performances in both approaches (Supplementary Fig. S7). Overall, for 1000 photons signal, and relatively in-plane molecules (*η* <60°), the orientation precision ranges stay below 15° for *ρ*, and below 37° for *δ*_2*D*_ (Supplementary Fig. S8).

**Figure 2:**
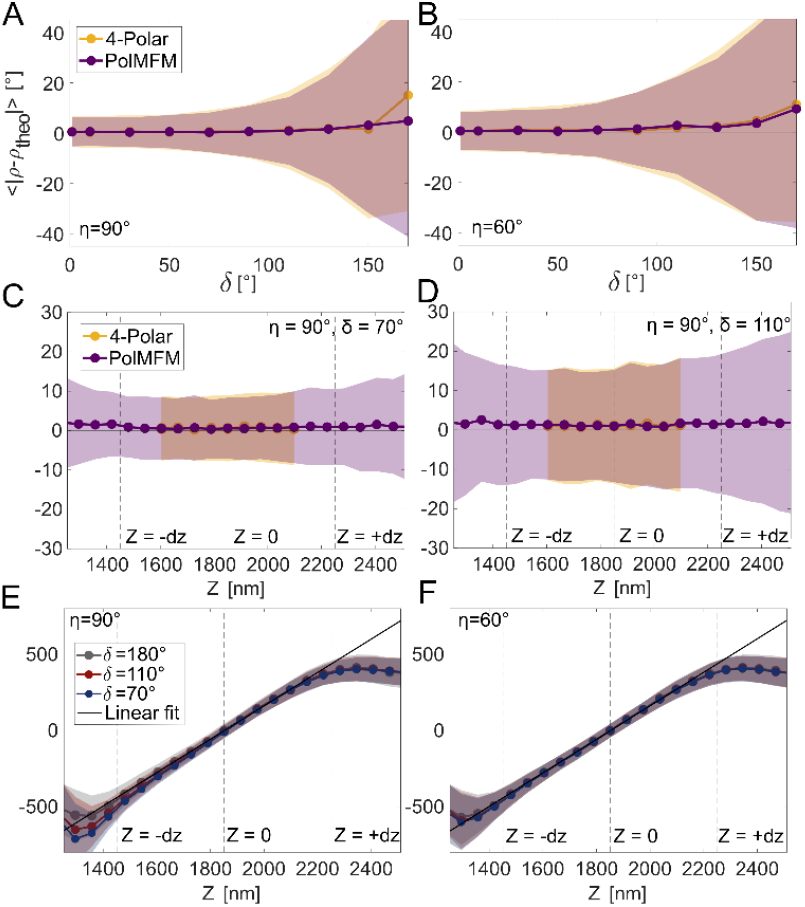
Bias and error on orientation estimation *ρ* and axial localization *z*_*GF*_ on simulated PSFs. For each simulated dipole of given orientation {*ρ, η, δ*}, 100 independent iterations of noise are generated. Each molecule emits 1000 photons and a background of 2 photons per pixel is added to the images. The graphs show the mean (dots) ± standard deviations (shaded areas) of the parameters retrieved over 100 iterations. A) Retrieved *ρ* as a function of wobbling *δ* for in-plane dipoles (η =90°) located axially at the central focal plane. Results are averaged for varying azimuthal orientation (ρ =0°: 15°: 165°). Two setups are simulated: the in-plane 4-Polar setup (yellow) and the PolMFM (purple). B) Same as A for inclined dipoles (η =60°). C) Retrieved ρ as a function of axial position *Z* for in-plane dipole (η =90°) with wobbling *δ* =70°. Results are averaged for varying azimuthal orientation (ρ =0°: 15°: 165°). D) Same as C with wobbling *δ* =110°. E) Retrieved axial localization *z*_*GF*_± standard deviation as a function of axial position of the dipole *Z* for in-plane dipoles (η =90°), with wobbling *δ* =180° (gray), *δ* =110° (red) or *δ* =70° (blue). Results are averaged for varying azimuthal orientation (ρ =0°: 15°: 165°). The retrieved values are fitted with a linear curve (black line) in the fully mobile case (*δ* =180°) which is taken as the ground truth (unaltered PSF shapes). F) Same as E for inclined dipoles (η =60°).

The impact of the axial position on orientation retrieval accuracy was next assessed. Molecules were simulated at various depths within a 1.2 µm axial range, laying in the sample plane (*η*= 90°) and exhibiting wobbling angles of *δ* =70° (Fig. 2C) and *δ* =110° (Fig. 2D). To mimic realistic experimental analysis pipeline, a localization filtering based on the standard deviation (sigma) of the fitted Gaussian(±30% of the theoretical sigma) was applied to exclude detections outside the depth of field. Despite splitting photons into six channels, PolMFM achieved comparable performance to the 4Polar scheme which splits photons into only four channels, while adding axial localization capabilities and providing orientation information across a larger depth of field, demonstrating its reliability for 2D orientation estimation. Finally, we examined the performance of the PolMFM in localizing single molecules in 3D by 3D Gaussian fitting. Fully flexible molecules (*δ* =180°) yielded the best localization accuracy due to their symmetric PSFs, while moderately constrained wobbling had only minor errors in the z-position estimation. Globally, the axial localization error ranged from 40 nm at the central focal plane of the PolMFM to 70 nm at the axial edge (Fig. 2E & F). We detected no axial localization bias across most of the axial range, except in the last 100 nm approaching the top focal plane, due to the dissymmetry of the information accessible from the 3 imaged planes. The results are comparable to previously reported performances for the 4- and 9-plane MFM configurations (^35,37^). Overall, these simulations confirm that PolMFM achieves reliable localization over nearly 1 µm axial range, a depth covered by the 3 focal planes.

Together, these simulations demonstrate the robustness of PolMFM in simultaneously retrieving molecular orientation and 3D position under realistic conditions, including noise, limited photon counts, variable wobble, and defocusing. Despite the polarization-sensitive defocused PSFs, PolMFM maintains high accuracy and precision.

### Experimental validation

To experimentally evaluate the performance of PolMFM, individual Janelia Fluor 549 molecules were immobilized in a polyvinyl alcohol (PVA) matrix and imaged under conditions allowing multiple consecutive frames detection per single molecule (Figs. 3A and S12-A). Spatiotemporally close detections were grouped (Material and Methods, *Molecular identification algorithm*) in order to quantify the orientation precision, defined as the standard deviation of repeated measurements for molecules detected in at least 10 frames. For a median photon count of 1000 photons (Supplementary Fig. S12-D) and a wobbling distribution centered at 90° (Supplementary Fig. S12-D), the in-plane orientation error distribution was centered around 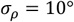, while the wobbling angle error centered around 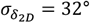 (Supplementary Fig. S12-F). These experimental values are consistent with simulations (Fig. 2 and Supplementary Fig. S7-8). As expected, the orientation precision degraded as the detected photon count decreased (Fig. 3B & C), ranging from 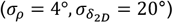 at 5000 photons to 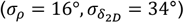 at 500 photons.

**Figure 3:**
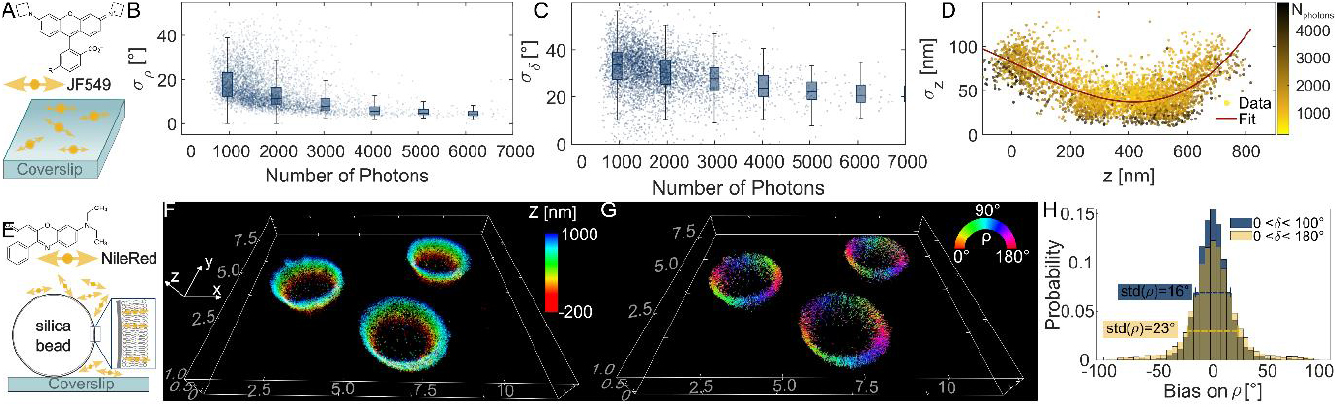
Experimental validation of PolMFM performance. A) Illustration of JF549 imaging embedded in PVA. B) Error on ρ as a function of number of detected photons. In the scatter plots, each point represents a molecule. In the box plots the statistics are computed for molecules withing a range of photon numbers. C) Same as B for wobbling retrieval *δ*_2*D*_. D) Axial localization error as a function of axial position, color-coded with mean photon number detected per molecule. A 3^rd^ order polynomial function (red curve) fits the error as a function of z. E) Illustration of NR imaging in a supported lipid bilayer covering a micrometric (3 µm) silica bead. F) 3D-SMLM reconstruction color-coded in z. Scale is in µm. G) 3D-SMOLM reconstruction, each localization is represented with a stick whose color and orientation correspond to retrieved in-plane angle *ρ*. The scale of the bounding box is in µm. H) Histograms of bias on orientation *ρ* − *ρ*_*theo*_. In blue, only dipoles exhibiting limited wobbling are kept (*δ* <100°), while in yellow dipoles exhibiting any wobbling are shown.

The localization precision was estimated as the standard deviation of the localizations from molecules detected between 10 and 30 times in consecutive frames. The lateral localization error distribution was centered at *σ*_*xy*_ =13 *nm* and the axial error at *σ*_*z*_=37 *nm* (Supplementary Fig. S12-B), in line with simulation results (in Figs. 2E & F and Figs. S9-A & C), and with previous results using MFM methods^35^. As expected, lateral localization precision improved with higher photon counts, reaching a median value of *σ*_*xy*_ =8 nm beyond 5000 detected photons (Supplementary Fig. S12-C). The axial localization precision on the other hand, depended on both the axial position of the molecule and on the photon count (Fig. 3D). Molecules axially located at the edge of the outer focal planes showed larger errors (*σ*_*z*_=70-90 nm depending on the signal level) compared to molecules near the central focal plane (*σ*_*z*_≈ 15-35 nm).

To evaluate the orientation retrieval accuracy in PolMFM on a more controlled system, we imaged Nile Red (NR), a hydrophobic dye soluble in lipids^42^, transiently bound to a supported lipid bilayer (SLB) coating silica beads of about 3 µm in diameter. The lipid composition, enriched in cholesterol (DPPC:Chol, 60:40), is known to reduce lateral diffusion and increase the membrane order, thereby influencing the orientation of Nile Red (NR) molecules^14^. Under these conditions, NR is expected to align perpendicular to the membrane surface (Fig. 3E) as previously shown^11,23,30^. The reconstructed 3D-SMLM (Fig. 3F) and SMOLM (Fig. 3G) images of the beads confirmed the capability of the PolMFM to efficiently retrieve 3D localizations and 2D orientations in realistic experimental conditions. The spherical geometry of the bead was used to calculate the expected in-plane orientation *ρ*_*theo*_ for each detected fluorophore based on its localization (Supplementary Note 5 and Supplementary Fig. S13), allowing to quantify the bias on orientation retrieval (*ρ* − *ρ*_*theo*_) for each detection (Fig. 3H). The resulting bias distribution is centered around 0°, demonstrating unbiased orientation retrieval over the whole PolMFM imaging volume, including for out-of-plane inclined dipoles. The standard deviation of the distribution was *σ*_*ρ*_ =23°, which is larger than in the JF549 dataset, despite restricting the analysis to molecules emitting more than 1500 photons (resulting in an average signal of about 2000 photons). This increased error likely results from a greater fluorophore wobbling, as indicated in the wobbling distribution centered around *δ*_2*D*_ ≈ 120° (Fig.S14-B). Simulations indicate that for such wobbling, the expected orientation error is between 17° and 23° for dipoles tilted moderately out-of-plane (*η*= 60° to 90°) (Fig 1A & B), which is consistent with the experimental findings.

To exclude dipoles with significant out-of-plane components (as proposed in^10^), analysis was restricted to molecules with limited projected wobbling (*δ*_2*D*_ <100°). This refinement reduced the orientation error to *σ*_*ρ*_ =16°, matching simulated values for in-plane dipoles (e.g. when *η* → 90°) while maintaining an unbiased distribution (Fig. 2A & C). Moreover, no dependence of the bias on the in-plane dipole orientation was observed (Supplementary Fig. S14-C). Together, these experiments confirm the PolMFM’s capacity to achieve precise and accurate 3D localization and 2D orientation measurements in SMLM experiments.

### 3D SMOLM imaging of actin filaments in fixed cells

Having validated PolMFM through simulations and test samples, we next applied the method to a biologically relevant sample—actin filaments—widely used as a benchmark in super-resolution and single-molecule orientation microscopy. Actin filaments, which are key components of the cellular cytoskeleton, can assemble into stress fibers with a well-defined nanoscale structure which has been used as a model system for the development of SMLM methods^43,44^. More recently, SMOLM techniques have enabled the measurement of the orientation of individual fluorophores labelling single actin filaments as well as more complex actin organizations in fixed cells such as stress fibers or the lamellipodia^9,10,30^.

Here, we used Alexa Fluor 568-phalloidin conjugates to label actin filaments in fixed U2OS cell as a reference sample to validate PolMFM (Material and Methods, *Actin imaging in fixed U2OS cells*) (Fig. 4). Phalloidin binds actin filaments with high affinity and provides a rigid anchor, enabling reliable orientation measurements^9^. The fluorophore’s molecular structure, together with its conjugation to phalloidin, ensures constrained rotational mobility suitable for SMOLM analysis^9,10,45^.

**Figure 4:**
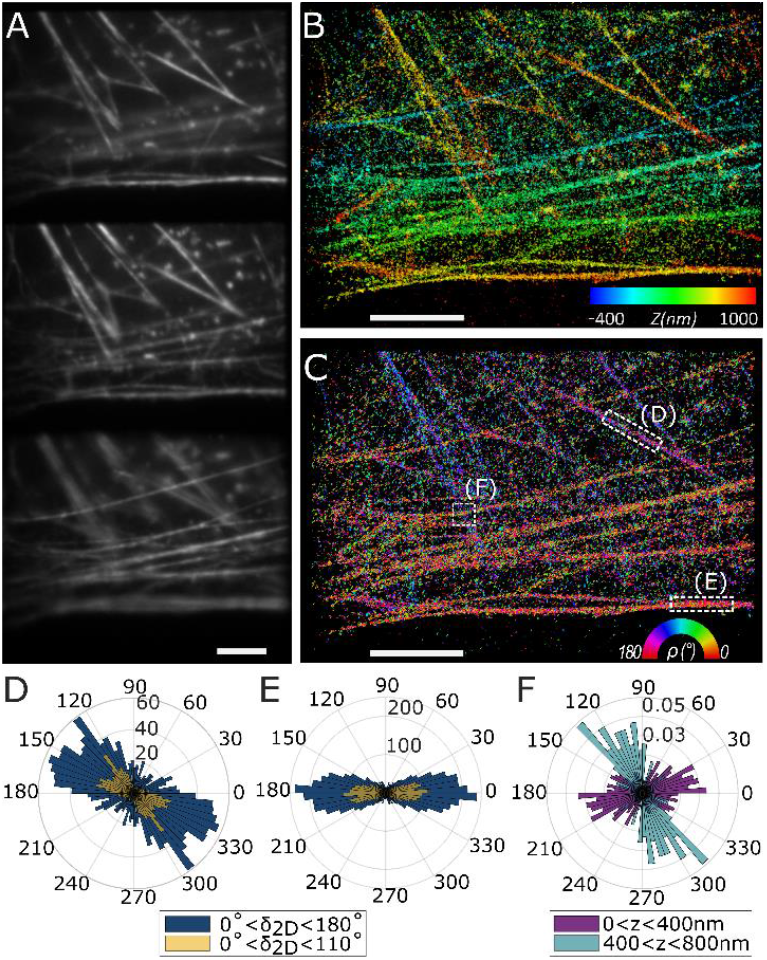
PolMFM imaging of actin filament organization in fixed U2OS cells labeled with AF568 phalloidin. A) Widefield 3D reconstructed image acquired with the PolMFM. The axial separation between each focal plane (from top to bottom) is Δ*z* = 370*nm*. B) 3D-SMLM reconstruction color-coded in z. C) SMOLM reconstruction, each localization is represented with a stick whose color and orientation correspond to retrieved in-plane angle *ρ*. Scale bar = 5µm. D) Polar histograms of *ρ* in the ROI (D) indicated by a white box in B), for all detections (blue) and detections exhibiting limited wobbling (yellow, *δ*_2*D*_ <110°). E) Same as D) for ROI (D). F) Polar histogram (probability) of *ρ* in the ROI (F), for all detections axially located between 0 and 400 nm (purple) and located between 400 and 800 nm (blue).

Figure 4 illustrates a PolMFM STORM acquisition in cells labeled with AF568-phalloidin. The three diffraction-limited images reconstructed from raw PolMFM widefield acquisition (Fig. 4A) reveal actin filaments at different heights, with distinct structures in focus in each plane. These structures are confirmed in the 3D-SMLM reconstruction, which exhibits fibers of different axial positions (Fig. 4B). Figure 4C displays the in-plane orientation estimated from PolMFM images, visualized as color-coded sticks representing the orientation angle. As expected, AF568 orientations align mainly along the main direction of the actin filaments. Histograms in selected regions confirm the reliable in-plane orientation retrieval, with standard deviations (*σ*_*ρ*_∼ 32° to 40°) (Fig. 4D–E) comparable to previous 4Polar STORM measurements in similar systems^10^ (*σ*_*ρ*_∼ 25° to 40°). Thresholding localizations with high wobbling reduces out-of-plane contributions and narrows the orientation distributions, improving precision for in-plane dipoles. Additional estimation of the error obtained on the parameter retrieval can be found in Supplementary Fig. S15, where measurements over multiple occurrences of single molecules lead to localization and orientation error in line with the modelling expectations described above.

Finally, Fig. 4F highlights a key advantage PolMFM over the conventional 4polar STORM method: the 3D axial position estimation of SMs, and thus the ability to separate filaments based on their axial position. As an illustrative example, the ROI analyzed in Fig. 4F contains two populations of different axial positions, shown in different colors. With traditional 4Polar, these populations would appear indistinguishable, as axial information is not available. In contrast, PolMFM is able to differentiate these filaments based on their axial localization, demonstrating its potential in resolving the orientation and organization of filament structures in three dimensions.

### 3D SMOLM imaging of DNA organization during spermiogenesis in the cricket

To demonstrate the potential of PolMFM in a biologically relevant and challenging context, we studied the DNA organization within the highly compact environment of sperm cell nuclei in the model cricket *Gryllus bimaculatus*. We focused on spermiogenesis, the process of cellular differentiation during which round spermatids undergo morphological and structural changes to become mature, motile sperm cells^46^. Accompanying these morphological changes, the sperm cells typically undergo a nuclear compaction involving dramatic volume reduction – often by an order of magnitude. In the model cricket *Gryllus bimaculatus*, the sperm nucleus adopts an elongated, needle-shaped morphology – common among many insects – with approximately 500 nm thick cross-section and ∼20 μm overall length, amounting to a ∼40-fold volume reduction compared to somatic nuclei.

In many animal species, this transition involves a dramatic genome-wide chromatin remodeling process^47^. In somatic and early germline cells, chromatin is organized as a basic unit called nucleosome, composed of an octamer of histone proteins around which ∼146bp of DNA is wrapped. During spermiogenesis, the vast majority of histones is removed and replaced by sperm-specific nuclear basis proteins (SNPBs), including protamines and protamine-like proteins. Protamines are highly basic proteins known to facilitate hyper-compaction. In vitro, protamines cause long DNA molecules (150kbp) to collapse into a toroidal organization^48^. In turn, short DNA molecules (<200bp) adopt nematic ordering and pack as discrete rod-like units in the presence of protamines^49^. These observations led to a model whereby “protaminized” DNA forms torus-like ordered structures in sperm^50^. Yet, how SNBPs organize DNA in vivo, and how this contributes to packaging the genome within the ultracompact sperm nucleus remains unclear^47,51^.

We and others previously showed that, in crickets, at an intermediate stage of nuclear elongation, referred to as canoe stage, electron microscopy (EM) images reveal that the genome organizes into ∼25 nm thick coiled chromatin fibers arranged as a stretched, highly regular spool along the nuclear axis^51^. This organization is thought to optimally compact the genome into the narrow, elongated needle-shaped nuclei of mature sperm cells. This highly regular organization in crickets is thus an ideal case-in-point to understand the precise arrangement of protamine-DNA complexes, but dedicated imaging technology is required to elucidate this dense volumetric organization within the 3D tissue.

Here, we extended prior EM, conventional fluorescence and SMLM (STORM) imaging description of genome reorganization during spermiogenesis by PolMFM study. We performed 3D SMOLM imaging of DNA fibers labelled with Sytox Orange, an intercalant DNA dye that orients perpendicular with respect to the DNA double-helix axis, in fixed cells. This orientation sensitivity enables inference of DNA fiber orientation based on fluorophore orientation^52,53^.

We dissected whole follicular testes from males, containing all intermediate stages of sperm maturation (Material and Methods, *Cricket testes fixed tissue preparation*). PolMFM imaging in fixed tissues across different stages enabled to track the evolution of DNA organization over the full nuclear volume at subsequent stages in spermiogenesis (Fig. 5).

**Figure 5:**
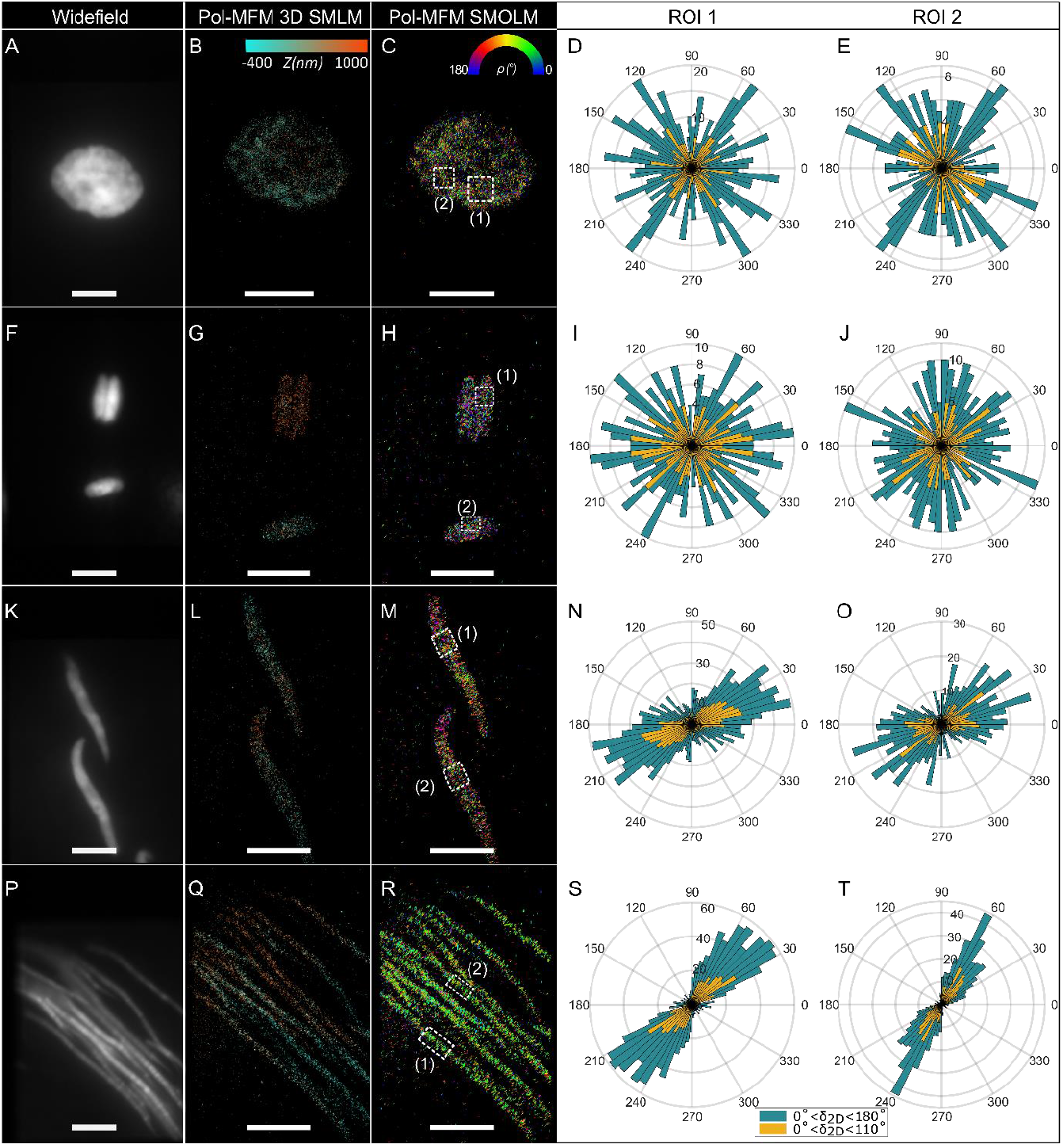
PolMFM imaging of Sytox-Orange labelling DNA in spermatid nucleus in cricket testes. Four different stages are identified: spermatocyte (A to E), early elongated spermatids (F to J), canoe spermatids (K to O) and needle-shaped spermatids (P to T). A-F-K-P) Widefield fluorescence image of the central focal plane. B-G-L-Q) 3D SMLM reconstruction color-coded for axial localization. C-H-M-R) SMOLM reconstruction where orientations *ρ* are shown as sticks and color coded for orientation. D-E-I-J-N-O-S-T) Polar histograms of orientations *ρ* in the corresponding ROIs at each stage. Blue histograms show all detections while yellow histograms are thresholded for the wobbling (*δ*_2*D*_ <110°). Scale bar = 5 µm.

As expected from the nucleosomal nature of chromatin in early stages, such as in spermatocytes (Fig. 5, A to C) and early elongated nuclei (Fig. 5, F to H), no preferential DNA orientation was identified. This is confirmed by the polar histograms of orientations computed for different ROI of the nuclei (Fig. 5D, E, I & J). At more advanced stages, when histones are no longer present in nuclei and thought to be replaced by SNBPs^51^, PolMFM imaging of canoe-shaped (Fig. 5, K to M) and needle-shaped (Fig. 5, P to R) nuclei, revealed strongly preferential orientations of Sytox Orange molecules. The polar histograms of orientations *ρ* (Fig. 5N, O, S & T), show that molecules tend preferentially to align perpendicular to the nuclei’s elongation axis. As the nucleus elongation progresses, the standard deviation of orientation distributions decreases from *σ*_*ρ*_ =36° - 37° in the canoe stage (Fig. 5N & O), to *σ*_*ρ*_ =26° - 30° in more advanced stages (Fig. 5T & S), suggesting a progressive increase of DNA fiber alignment.

Finally, we combined results from multiple cells at each stage to quantify the narrowing of orientation distributions. Histograms of *ρ* revealed a progressive narrowing from the canoe stage onward, indicating a progressive alignment of DNA molecules within spooled fibers during spermiogenesis (Supplementary Fig. S16).

These results are consistent with previous EM studies reporting elongated, axially aligned fibers at late spermiogenesis^51^. While EM was not able to directly resolve the underlying DNA organization within the fibers, PolMFM offers a direct means to reveal the molecular ordering in such fibers. Moreover, the access to a large axial range in the tissue (Fig. 5) permits to report its DNA complex organization under volumetric conditions, which is made possible by using highly inclined illumination (Material and Methods, *Cricket testes fixed tissue preparation*). Our findings support the hypothesis that sperm chromatin fibers are composed of discrete bundles of nematically ordered DNA molecules in the elongated spermatid nucleus of the cricket, and that this elongation is achieved progressively through the spermiogenesis process.

## Discussion

In this work, we introduced PolMFM as a new tool for extracting both in-plane orientation and rotational wobbling of single molecules, in addition to their 3D localization. PolMFM combines images from three focal planes and four polarization projections, without modifying notably the single molecules’ point spread function (PSF) shape. PolMFM’s design allows straightforward orientation and localization retrieval, using intensity analysis across the polarization-projection channels^10,30^ and standard 3D Gaussian fitting ^35,36^.

A central question in polarization-resolved microscopy is whether defocusing affects the interpretability of intensity projections across polarization channels. We show that defocusing has a minimal impact on the accuracy of orientation estimates, essentially because defocusing spreads the signal over a larger area but does not alter the intensity distribution across polarization channels. Similarly, due to the nature of PolMFM processing based on integrated intensities, we also expect this method to be insensitive to geometrical aberrations, which is a strong advantage as compared to analyzes based on the PSF shapes.

Our simulations confirmed that projecting polarized PSFs over multiple focal planes does not degrade orientation estimation. Importantly, both the accuracy and precision of orientation retrievals remain comparable to those of in-plane 4-polarization schemes throughout the full depth-of-field of PolMFM. Likewise, 3D localization accuracy is preserved, as signals from orthogonal polarization channels are recombined to reconstruct an unpolarized 3D PSF.

Our simulations also define the conditions under which PolMFM maintains robust orientation and localization estimations. Across the full axial detection range, determining the orientation and localization remain accurate and precise for molecules exhibiting moderate wobbling (*δ* ≈ 60°–120°). For highly flexible dipoles, orientation is inherently undefined. Conversely, for rigid dipoles, the PSF is strongly deformed^40,41^. In this case, accurate localization would require more complex PSF models that account for dipole orientation and axial position, as used in PSF engineering approaches. This would significantly increase computational complexity of our method. Nevertheless, it has been found in many applications that the labeling conditions produce moderate wobbling (*δ* ≈80°–120°), allowing PolMFM to retrieve unbiased localization while reporting orientation, thus achieving robust 3D localization and 2D orientation estimation without complex PSF modeling.

Owing to its compatibility with routine dense SMLM acquisition, we demonstrated that PolMFM enables the simultaneous retrieval of both orientations and 3D localizations of AF568-phalloidin–labeled actin filaments in fixed mammalian cells, with improved 3D imaging capabilities over a 1.2 µm axial range compared to the conventional 4-polarization splitting scheme^10,30^.

Finally, we applied PolMFM to decipher the progressive spatial reorganization of DNA during spermiogenesis in the cricket model system of *Grillus bimaculatus*, a challenging, biologically relevant context due to the dense and volumetric nature of sperm nuclei. PolMFM enabled widefield, in-tissue SMOLM imaging, capturing molecular orientation several microns deep within testis tissue. This approach extends beyond previously developed SMOLM methods, which generally use total internal reflection fluorescence (TIRF) illumination that limits the studies to thin samples. PolMFM could thus provide orientation data across the entire nuclear volume, complementing previous EM and fluorescence studies on DNA-SNBP arrangements^51^.

Many SMOLM methods have been developed over the past years, some enabling the full 6D localization and orientation parameters determination^11,13,14,22–24,26,54^. Most methods require complex PSF modelling and fitting, are sensitive to calibration and aberrations, and are often not well-suited for imaging dense sample due to their extended PSF spatial footprint^17^. The raMVR method^11^ stands out for its excellent performance in terms of measurement precision and robustness to aberrations over a depth range of 1.5µm, while maintaining a relatively compact PSF. However, this performance comes at the cost of complex optical design that is challenging to implement and align. PolMFM offers an accessible and robust alternative for applications where the access to 2D orientation is sufficient, decreasing the complexity of both instrumentation and data analysis.

While MFM requires the design of custom diffractive gratings and prisms, simpler implementations using beam splitters are possible^55–57^, thus making the method accessible to a broader range of laboratories. In particular, a 4-Polar STORM setup can be upgraded by introducing a defocus between two of its four polarization channels, enabling 3D localization in addition to the extraction of molecular orientation and wobbling. Such an implementation would come at the cost of reduced axial localization range compared to PolMFM. Conversely, the axial range could be expanded using a four-plane MFM before polarization splitting, yielding eight images. Such configurations are compatible with SMLM, as 9-plane MFM has already been demonstrated^35^.

Finally, PolMFM could be extended to ensemble polarization analysis to enable diffraction-limited organization imaging. Provided appropriate analysis, this implementation would allow to build instantaneous 3D orientation-maps of biological samples, as proposed previously using sequential polarization excitation^58^, prior to molecular-scale organization investigation.

In conclusion, PolMFM provides a robust, accessible method for simultaneous 3D localization and in-plane orientation measurement of single molecules, without relying on complex PSF engineering or computationally intensive reconstruction. Its compatibility with dense SMLM imaging and in-tissue acquisition makes it a powerful tool for probing molecular organization in 3D across various biological contexts, from membranes to cytoskeleton to chromatin.

## Materials and Methods

### PolMFM Optical Setup

The PolMFM setup is integrated into a conventional wide-field fluorescence microscope (Nikon Eclipse Ti2). Excitation is provided by a set of continuous-wave lasers emitting at 640nm (LPX-640L, OXXIUS), 560 nm (Cobolt Jive 200 561nm), 488nm (LBX-488-100, OXXIUS) and 405nm (Cobolt MLD 405nm). The laser lines are merged into a single path, and an acousto-optical tunable filter (AOTFnC-400.650-TN, AA Opto-Electronic) is introduced to modulate independently their relative intensities. An additional custom-made electronic circuit, incorporating an Arduino Due board and electronic switches, was used to modulate the AOTF and to synchronize the laser excitation to the camera exposure. The excitation beam is expanded using a telescope before entering the microscope. A quarter-wave plate (QWP, AQWP10M-580 Thorlabs) is introduced in the path of the linearly polarized light, to achieve near-circular polarization excitation. The beam is focused in the back focal plane of the objective using a 500mm lens (f = 500 mm, ACT508-500-A Thorlabs), creating an illumination field of view of approximately 17 µm in diameter (FWHM). The 500 mm lens, quarter-wave plate (QWP), and mirror were mounted on a linear translation stage to laterally displace the focused beam in the back focal plane (BFP) without introducing tilt, enabling manual switching between EPI and highly inclined and laminated optical sheet (HILO) illumination modes. The excitation light is reflected on a dichroic mirror (Di03-R405/488/561/635, ref. FL-007139, Semrock) and directed to the sample via an oil immersion objective lens (Plan Apo 100X λD, OI, NA = 1.45, Nikon). The objective lens is mounted on a piezo electric stage to allow for precise control of the z-axis displacement (PIFOC, Physik Instrumente).

Fluorescence emitted by the sample is collected by the same objective lens and transmitted through the dichroic mirror. At the exit of the microscope, the emitted light passes through a second but identical dichroic mirror (FL-007139, Semrock), positioned with an inverted plane of incidence to compensate the polarization distortions introduced by the first dichroic mirror. A slit (SP 60, OWIS) positioned at the microscope’s primary image plane delimits the final field of view (∼20 × 25 µm^2^) on the camera and avoids any overlap between the different projection channels. A diffraction multi-focus grating (MFG, custom made by Silios) is placed on the emission path of the microscope in conjugation to the back focal plane of the objective lens thanks to a relay lens (f_1_ = 150 mm, AC508-150-A Thorlabs). The specific design of the MFG splits the light into three diffraction orders with equivalent intensity reparation, each corresponding to a different focal plane, forming three spatially separated images on the camera (Fig. 1A). A chromatic correction grating (CCG) made of a blazed grating followed by a multi-faceted prism are used to correct for chromatic dispersion and to tilt the 3 diffractive orders toward the camera. A half-wave plate (½-inch diameter, 05RP02-24, Newport) is placed after the prism and positioned to selectively cover the central focal plane and rotate its polarization by 45°. A Wollaston prism with a circular aperture of 30 mm and a splitting angle of 1° (PWQ60.30, Bernhard Halle Nachfl. GmbH) is introduced before the refocusing lens (f_2_ = 200 mm, ACT508-200-A Thorlabs). The Wollaston is oriented to project vertically the outer focal planes into 0° and 90° polarizations, and the central focal plane into 45° and 135° polarizations. Finally, a band-pass filter (FF01-617/73-25, Semrock) is positioned before the EMCCD imaging camera (DU-897U-CSO-#BV, iXon Ultra, Andor). The effective pixel size on the camera is 120 nm.

Focus stability during measurements is maintained using a commercial autofocus system (Perfect Focus System, Nikon). The acquisition hardware is controlled using a customized commercial software (Inscoper).

Before each acquisition round, several calibration procedures ensuring channel registration, intensity imbalance correction and polarization calibration, are applied. The procedures are detailed in SI Note 3.

### Simulations

The aim is to simulate the emission of a fluorescent molecule and its propagation through an optical system, both the PolMFM and the conventional in-plane 4-Polar setup^10^. We model a fluorescent molecule with an oscillating electric dipole 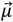, and test multiple wobbling values *δ*, multiple angular parameters {*ρ, η*} and multiple localizations {*x, y, z*}. The image of a molecule rotating within a wobbling cone on timescale faster than the exposure time of the camera is identical to the image produced by three fixed orthogonal super-imposed dipoles 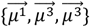, of amplitudes 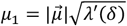 and 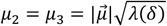, where *λ*(*δ*) is given by equation S9 in Supplementary Note 1.2^39^. After propagation through the optical system, we can write the polarized electric fields in the different channels of the PolMFM as follows:

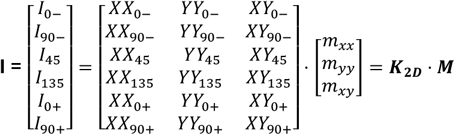

where 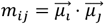 ({*i, j*} ∈ {*x, y*}) are the second-order dipole moments, and the coefficients of the propagation matrix ***K***_**2D**_ are 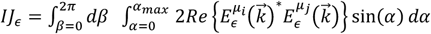 with *I, J* ∈ *X, Y*} and *ϵ* is the direction of polarization projection (*ϵ* ∈ {0, 90, 45, 135}). The coefficients of ***K***_**2D**_ account for the propagation through the microscope, with integration over all propagation directions 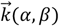 (Supplementary Note 1.3 and Supplementary Fig. S2). (*α, β*) represent the coordinates in the BFP and account for the collection angle and in-plane polar-angle respectively. *α*_*max*_ is the numerical aperture of the microscope. The electric field 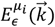 polarized along *ϵ* emitted by a fixed dipole 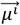 is given by a vectorial model of light propagation through a refractive index mismatch interface, developed in^59^ (Supplementary Note 2). The microscope parameters for the simulations are specified in Table S1. To achieve realistic image simulations, camera and background noises were incorporated. The camera noise model proposed in ^10^ was applied, given the similarity in hardware and experimental conditions. Simulations also allowed for varying background levels. Each simulated molecule emitted a total of 1,000 photons, distributed across the 6 PolMFM channels, with a constant background of 2 photons per pixel. For every combination of orientation parameters (*ρ, η, δ*) and axial position *z*, 100 independent noise realizations were generated to perform Monte Carlo simulations. In the case of the 4-Polar scheme, the total number of photons was split into only 4 channels. The bias was determined as the difference between the simulated theoretical value and the retrieved value averaged over the multiple noise iterations. The error was defined as the standard deviation over multiple iterations of the same noisy measurement.

To retrieve the localization parameters {*x*_0_, *y*_0_, *z*_0_}, the unpolarized 3D PSFs are first built from the 6 polarized channels of the PolMFM by summing the two perpendicularly polarized images in each focal plane. A 3D-Gaussian fitting is performed on these 3D PSFs to retrieve the localization of the molecule, as done in the analysis pipeline (see Fig 1C-3). Then the intensities detected in each channel ***I*** =[*I*_0−_ *I*_90−_ *I*_45_ *I*_135_*I*_0+_ *I*_90+_]^*T*^are computed by summing the pixel intensities inside a box centered around each PSF, where the local background has been removed:

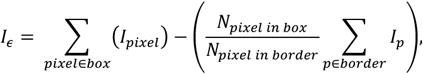

where the size of the box is defined by a half box extension parameter *I*_*box*_ =5 pixels and the full box extension is *box* =[*x*_*GF*_− *I*_*box*_ ∶ *x*_*GF*_+ *I*_*box*_], “*border*” corresponds to the 1 pixel large outer border of the box, *N*_*Pixel in box*_ is the number of pixels contained in “*box*” and *N*_*pixeI in border*_ is the number of pixels contained in “*border*”. The second-order dipolar moments are inferred from the operation ***M*** =(***K***_**2D**−***theo***_)^−1^ ⋅ ***I***, where the theoretical propagation matrix is:

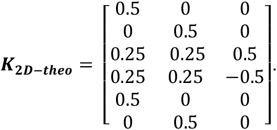

Finally, the orientation parameters {*ρ, δ*_2*D*_} are computed using Supplementary Equation S19. While ***K***_**2D**−***theo***_ is used in the context of the Monte-Carlo simulations, the experimental propagation matrix is inferred from a polarization calibration step (Supplementary Note 4). The experimental matrix ***K***_**2D**_ thus incorporates polarization aberrations coming from the setup. An example of experimental matrix is given below:

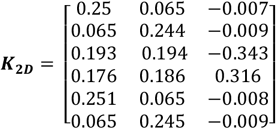

### JF549 in PVA matrix

A 5% w/w PVA solution was prepared by dissolving 1.25 g of PVA powder (molecular weight 146,000-186,000, ref. 363103, Sigma Aldrich) in 25 mL of Milli-Q water. The dissolution was carried out in a round-bottom flask placed in a silicon oil hot bath at 96°C under gentle stirring and reflux for 1h. The solution was sealed and stored at 4°C. JF549-Halo dyes were diluted to 1 nM final concentration in the 5% PVA solution. 500 µL of the solution was deposited onto a cleaned coverslip and spin-coated for 30 s at 500 rpm, followed by 30 s at 1000 rpm. The coated coverslips were polymerized by incubation at 70°C overnight. To achieve single-molecule regime, the 560 nm laser intensity is adjusted to reach 22 kW/cm^2^ at the sample. We acquired between 2000 and 7000 frames in several fields of view with an exposure of 50 ms and a camera gain of 150.

### NR-labelled SLB-coated beads

#### Preparation of the SLB-coated beads

The preparation is based on the protocol proposed in^30^. First, silica beads of 3 µm diameter (Kisker Biotech GmbH & Co.) were diluted to 2mg/mL as follows. The bead stock solution (at 50 mg/mL) was vortexed for 10 seconds and centrifuged for 5 minutes at 14500 rpm. After removing the supernatant liquid, the beads were resuspended in a TRIS buffer (20 mM TRIS and 50 mM KCl, pH 7.5). After 3 cycles of vortex, centrifugation and resuspension, the bead solution was put in a water bath at 70°C for 15 minutes, then supplemented with 20 mM of CaCl_2_, vortexed for 30 seconds and kept in the water bath at 70°C for 15 additional minutes. Second, the lipid solution is prepared as follows. DPPC (60%) and cholesterol (40%) (Avanti Polar Lipids), were dissolved in chloroform (VWR chemicals). The chloroform was evaporated under a gentle flow of nitrogen gas, resulting in a thin lipid film on the vial that was kept overnight in a vacuum chamber to remove any excess residue of organic solvent. The lipid film was then rehydrated in the TRIS buffer (pre-warmed at 70°C) to a concentration of 1.4 mg/mL. The lipid suspension was incubated for 30 minutes at 70°C and vortexed every 5 minutes. Finally, the hydrated lipid film was sonicated for 30 minutes in a bath 70°C until the solution became completely transparent, indicating the formation of small vesicles. Finally, to coat the beads with lipid bilayers, the two solutions (lipid and bead solutions) were mixed in a volume ratio of 1:1 and vortexed. The mixture was incubated in a water bath at 70°C for 1 hour, while being vortexed every 10 minutes to allow the vesicle to fuse with each other and resulting in a complete coating of the beads with lipid. The lipid bilayer-coated beads were then allowed to cool gradually to an ambient temperature for an hour, while vortexed every 15 minutes. To remove excess vesicles from the coated bead solution, the solution was centrifuged at 5000 rpm for 5 minutes at room temperature (RT). The supernatant was trashed, and the pellet was replenished with the TRIS buffer and vortexed. This procedure was repeated 5 times. The lipid-coated beads were stored at 4°C for a maximum of 2 weeks for experiments.

#### Preparation of the SLB-coated beads with Nile Red labelling for imaging

The coverslips (µ-Slide 8 Well Glass Bottom, Ibidi, ref. 80827) were coated with 0.01% poly-L-lysine (Sigma, ref. P4832) for 1.5 hours, followed by three washes in Tris buffer. One volume of the SLB-coated bead solution was vortexed prior to use and diluted with three volumes of Tris Buffer. Nile Red was added to the mixture at a final concentration of 750 nM. The resulting solution was added to the wells for imaging.

#### Acquisition parameters

Prior to the SM acquisition, a widefield fluorescence z-stack was acquired using a 100 nm axial step and low 560 nm laser power (∼10 to 40 W/cm^2^). Then, the laser power is raised to ∼22 kW/cm^2^ for SM imaging. For each field of view, between 30,000 and 40,000 frames were recorded at a frame rate of 20-30 ms, with the camera electronic gain set to 250.

### Actin imaging in fixed U2OS cells

#### Cell culture

Experiments were carried out on U2OS (human osteosarcoma) wildtype cells. U2OS cells were grown in an incubator at 37°C and 5% CO2 in DMEM (Gibco, ref. 31053028) supplemented with 10% fetal bovine serum (FBS, Dutscher, ref. S1900-500C), 1% Glutamax (Gibco, ref. 35050061) and 1% Penicillin-Streptavidin (Gibco, ref. 15140-122). Cells were passaged every 2 to 4 days with trypsin (Gibco, ref. 12605010) and maintained at a density ranging from 0.2 · 10^5^ to 1.1· 10^5^ cells · cm^−1^. Cells were checked for mycoplasma infection every 3-4 months and tested negative.

#### Cell preparation for actin imaging

Coverslips (#1 thickness) were cleaned by sequential sonication for 5 min each in acetone, ethanol, and Milli-Q water, followed by 5 min of air plasma cleaning. The cleaned coverslips were coated with 10 µg/mL fibronectin (Bovine Fibronectin, Sigma-Aldrich, ref. F1141) for 2 hours at RT. U2OS cells were seeded at a density of ∼2.4 · 10^5^ cells · cm^−2^. For fixation, cells were incubated with 4% paraformaldehyde (PFA) (EMS, ref. 15710) in cytoskeleton buffer (10 mM MES, 150 mM NaCl, 5mM EGTA, 5mM Glucose, 5 mM MgCl_2_, pH 6.1) for 15 minutes at RT. Fixed cells were washed three times for 5 min each with PBS. Permeabilization and blocking were performed using P/B buffer (10% BSA, 0.1% Saponin in PBS) for 2.5 hours at RT. Cells were labelled with Alexa Fluor 568 (AF568)-phalloidin (Invitrogen, ref. A12380) by incubating overnight at 4°C in P/B buffer containing ∼ 300 nM dye. Cells were washed twice for 5 min with PBS prior to imaging.

#### PolMFM imaging in STORM buffer

For PolMFM experiments, coverslips were mounted in AttoFluor cell chambers (Invitrogen, ref. A7816) with STORM imaging buffer (50mM Tris at pH=8, 10 mM NaCl, 10% glucose, 100 mM MEA, 56 U/mL Glucose Oxidase, 340 U/mL Catalase, prepared as described in ^60^). The sample was covered by a glass coverslip to minimize oxygen exchanges. The widefield image was obtained by averaging 10 frames acquired with a 560 nm laser power of 50 W/cm^2^. For the SM acquisition, 15000 frames were acquired in epi-illumination mode, with an exposure time of 50ms at a laser power of ∼15 kW/cm^2^ and the camera EM gain was set to 200.

### Cricket testes fixed tissue preparation

#### Rearing conditions

*Gryllus bimaculatus* were reared at 29°C with a 12h:12h light:dark cycle and fed with standard meal powder (see ^51^).

#### Testicular follicle preparation

*H*ealthy >7 days old male adults were dissected in PBS containing 0,15% Triton (PBS-T) medium, testes were dilacerated, rinsed in PBS and fixed in 4% PFA in PBS solution for 10 min. Testes were transferred to a PBS solution containing 50% acetic acid for 3 min on a coverslip, then squashed manually with a polylysine-coated glass slide. Squashed testes were dipped in liquid nitrogen, the coverslip was quickly removed, and slides were transferred to 100% ethanol bath at −20° C for at least 10min and up to several weeks. Before imaging, slides were rehydrated by incubation in three successive PBS baths at RT. Sample were stained and permeabilized with 500 nM Sytox Orange (Invitrogen, ref. S11368) and 0.5% Triton X-100 (Sigma Aldrich, ref. T8787) in PBS, then mounted and sealed with coverglass without washing.

#### PolMFM imaging

Imaged cells were localized at different depths in the tissue, reaching a few tens of µm away from the coverslip. We used HILO (Highly Inclined and Laminated Optical sheet) excitation to reduce background arising from out-of-plane tissue contributions. For each field of view, between 10,000 and 20,000 frames were acquired with an exposure time of 40-50ms and a camera gain of 200. The molecular blinking was achieved with a 560 nm laser power ranging from 19 and 22 kW/cm^2^, and progressive 405 nm laser activation up to 60 W/cm^2^.

### Molecular identification algorithm

The molecular identification algorithm used in this work to correct for molecular blinking is based only on spatial and temporal thresholds and was previously introduced ^61^. The code iterates over all the frames of the input point-cloud dataset. For each localization in a given frame, distances to all localizations within a user-defined temporal window (e.g. over 20 frames if the search time is defined at 1 second and the acquisition frame rate is 50ms) are computed to account for possible reversible dark states. Localizations that appear in the following frames within the defined time window and that are separated by less than a user-defined spatial threshold are grouped and assigned a common molecular identifier.

The algorithm returns an index for each localization corresponding to a given unique molecule identifier. Based on this information, a mean position and a mean frame number are computed for each of the multiple-appearing molecules. This process generates a new point cloud data set with a different number of elements compared to the original one, reflecting the consolidation of multiple localizations into single molecular events.

In the present work, the output table was expanded to include additional metrics for each identified molecule, such as the standard deviation of the position, mean in-plane orientation angle (*ρ*) and wobbling angle (*δ*), along with their respective standard deviations, and the mean photon count. This algorithm was applied on the data acquired with JF549 in PVA, with the following parameters: Δ*x* = 30 *nm* and Δ*z* = 100 *nm* for the spatial threshold, and Δ^T^ = 250 *ms* for the temporal threshold.

## Acknowledgments

BH acknowledges funding from, the Agence National de la recherche (ANR-19-CE42-0003-01), the LabEx CelTisPhyBio (ANR-11-LABX-0038, ANR-10-IDEX-0001-02), the Institut Curie, DIM ELICIT. BH and GO acknowledge ITMO Cancer of Aviesan on funds administered by Inserm (grant N° 20CP092-00). BH and SB recognizes the support of France-BioImaging infrastructure grant ANR-10-INBS-04 (Investments for the future). LR acknowledges a doctoral grant from École Doctorale Physique en Île-de-France (EDPIF). S.D. and F.-C.T. are funded by ANR grant ANR-20-CE11-0010-01 (ActinFission) and PSL-Qlife grant (MyoMemActin). S.D. is funded by a post-doc fellowship from Fondation ARC.

## Author Contributions

G.A.O., S.B. and B.H. designed the study. L.R. and B.H. designed and mounted the microscope. L.R. programmed and performed simulations. L.R., G.A.O., S.D. and F.T.C. prepared the samples. L.R., G.A.O. and C.V.R acquired the data. L.R. and B.H. developed the analysis pipeline. L.R. analyzed the data. L.R., G.A.O., S.B. and B.H. prepared the manuscript. All authors contributed to the writing of the paper.

## References

1. Shroder, D. Y., Lippert, L. G. & Goldman, Y. E. Single molecule optical measurements of orientation and rotations of biological macromolecules. Methods Appl Fluoresc 4, 042004 (2016).

2. Zhang, O. & Lew, M. D. Single-molecule orientation localization microscopy II: a performance comparison. J. Opt. Soc. Am. A, JOSAA 38, 288–297 (2021).

3. Alonso, M. & Brasselet, S. Polarization microscopy: from ensemble structural imaging to single molecule 3D orientation and localization microscopy. Optica (2023) doi:10.1364/OPTICA.502119.

4. Zhang, O. & Lew, M. D. Single-molecule orientation-localization microscopy: Applications and approaches. Quarterly Reviews of Biophysics 57, e17 (2024).

5. Sharonov, A. & Hochstrasser, R. M. Wide-field subdiffraction imaging by accumulated binding of diffusing probes. Proceedings of the National Academy of Sciences 103, 18911–18916 (2006).

6. Betzig, E. et al. Imaging Intracellular Fluorescent Proteins at Nanometer Resolution. Science 313, 1642–1645 (2006).

7. Hess, S. T., Girirajan, T. P. K. & Mason, M. D. Ultra-high resolution imaging by fluorescence photoactivation localization microscopy. Biophys J 91, 4258–4272 (2006).

8. Rust, M. J., Bates, M. & Zhuang, X. Sub-diffraction-limit imaging by stochastic optical reconstruction microscopy (STORM). Nat Methods 3, 793–796 (2006).

9. Valades Cruz, C. A. et al. Quantitative nanoscale imaging of orientational order in biological filaments by polarized superresolution microscopy. Proceedings of the National Academy of Sciences 113, E820–E828 (2016).

10. Rimoli, C. V., Valades-Cruz, C. A., Curcio, V., Mavrakis, M. & Brasselet, S. 4polar-STORM polarized super-resolution imaging of actin filament organization in cells. Nat Commun 13, 301 (2022).

11. Zhang, O. et al. Six-dimensional single-molecule imaging with isotropic resolution using a multi-view reflector microscope. Nat. Photon. 17, 179–186 (2023).

12. Zhang, O., Zhou, W., Lu, J., Wu, T. & Lew, M. D. Resolving the Three-Dimensional Rotational and Translational Dynamics of Single Molecules Using Radially and Azimuthally Polarized Fluorescence. Nano Lett 22, 1024–1031 (2022).

13. Ding, T. & Lew, M. D. Single-Molecule Localization Microscopy of 3D Orientation and Anisotropic Wobble Using a Polarized Vortex Point Spread Function. J. Phys. Chem. B 125, 12718–12729 (2021).

14. Lu, J., Mazidi, H., Ding, T., Zhang, O. & Lew, M. D. Single-Molecule 3D Orientation Imaging Reveals Nanoscale Compositional Heterogeneity in Lipid Membranes. Angewandte Chemie International Edition 59, 17572–17579 (2020).

15. Hecht, E. Optics. (Pearson, Boston Columbus Indianapolis New York San Francisco Amsterdam Cape Town Dubai London Madrid Milan Munich, 2017).

16. Alonso, M. & Brasselet, S. Polarization microscopy: from ensemble structural imaging to single molecule 3D orientation and localization microscopy. Optica (2023) doi:10.1364/OPTICA.502119.

17. Zhang, O. & Lew, M. D. Single-molecule orientation-localization microscopy: Applications and approaches. Quarterly Reviews of Biophysics 57, e17 (2024).

18. Brasselet, S. & Lew, M. D. Single-molecule orientation and localization microscopy. Nat. Photon. 19, 925–937 (2025).

19. Bartko, A. P. & Dickson, R. M. Imaging Three-Dimensional Single Molecule Orientations. J. Phys. Chem. B 103, 11237–11241 (1999).

20. Böhmer, M. & Enderlein, J. Orientation imaging of single molecules by wide-field epifluorescence microscopy. J. Opt. Soc. Am. B, JOSAB 20, 554–559 (2003).

21. Toprak, E. et al. Defocused orientation and position imaging (DOPI) of myosin V. Proceedings of the National Academy of Sciences 103, 6495–6499 (2006).

22. Hulleman, C. N. et al. Simultaneous orientation and 3D localization microscopy with a Vortex point spread function. Nat Commun 12, 5934 (2021).

23. Wu, T., Lu, J. & Lew, M. D. Dipole-spread-function engineering for simultaneously measuring the 3D orientations and 3D positions of fluorescent molecules. Optica, OPTICA 9, 505–511 (2022).

24. Curcio, V., Alemán-Castañeda, L. A., Brown, T. G., Brasselet, S. & Alonso, M. A. Birefringent Fourier filtering for single molecule coordinate and height super-resolution imaging with dithering and orientation. Nat Commun 11, 5307 (2020).

25. Wu, T., Lu, P., Rahman, M. A., Li, X. & Lew, M. D. Deep-SMOLM: deep learning resolves the 3D orientations and 2D positions of overlapping single molecules with optimal nanoscale resolution. Opt. Express, OE 30, 36761–36773 (2022).

26. Jouchet, P., Roy, A. R. & Moerner, W. E. Combining deep learning approaches and point spread function engineering for simultaneous 3D position and 3D orientation measurements of fluorescent single molecules. Optics Communications 542, 129589 (2023).

27. Backer, A. S., Lee, M. Y. & Moerner, W. E. Enhanced DNA imaging using super-resolution microscopy and simultaneous single-molecule orientation measurements. Optica, OPTICA 3, 659–666 (2016).

28. Ohmachi, M. et al. Fluorescence microscopy for simultaneous observation of 3D orientation and movement and its application to quantum rod-tagged myosin V. Proceedings of the National Academy of Sciences 109, 5294–5298 (2012).

29. Mehta, S. B. et al. Dissection of molecular assembly dynamics by tracking orientation and position of single molecules in live cells. Proceedings of the National Academy of Sciences 113, E6352–E6361 (2016).

30. Bruggeman, E. et al. POLCAM: instant molecular orientation microscopy for the life sciences. Nat Methods 21, 1873–1883 (2024).

31. Kumar, C. S. S. et al. 4polar3D: Single molecule 3D orientation imaging of dense actin networks using ratiometric polarization splitting. 2025.07.13.664601 Preprint at 10.1101/2025.07.13.664601 (2025).

32. Huang, B., Wang, W., Bates, M. & Zhuang, X. Three-Dimensional Super-Resolution Imaging by Stochastic Optical Reconstruction Microscopy. Science 319, 810–813 (2008).

33. Pavani, S. R. P. et al. Three-dimensional, single-molecule fluorescence imaging beyond the diffraction limit by using a double-helix point spread function. Proceedings of the National Academy of Sciences 106, 2995–2999 (2009).

34. Shechtman, Y., Weiss, L. E., Backer, A. S., Sahl, S. J. & Moerner, W. E. Precise Three-Dimensional Scan-Free Multiple-Particle Tracking over Large Axial Ranges with Tetrapod Point Spread Functions. Nano Lett. 15, 4194–4199 (2015).

35. Hajj, B. et al. Whole-cell, multicolor superresolution imaging using volumetric multifocus microscopy. Proc. Natl. Acad. Sci. U.S.A. 111, 17480–17485 (2014).

36. Abrahamsson, S. et al. Fast multicolor 3D imaging using aberration-corrected multifocus microscopy. Nat Methods 10, 60–63 (2013).

37. Hajj, B., Oudjedi, L., Fiche, J.-B., Dahan, M. & Nollmann, M. Highly efficient multicolor multifocus microscopy by optimal design of diffraction binary gratings. Sci Rep 7, 5284 (2017).

38. Hecht, E. Optics. (Pearson, Boston Columbus Indianapolis New York San Francisco Amsterdam Cape Town Dubai London Madrid Milan Munich, 2017).

39. Backer, A. S. & Moerner, W. E. Determining the rotational mobility of a single molecule from a single image: a numerical study. Opt. Express, OE 23, 4255–4276 (2015).

40. Lew, M. D., Backlund, M. P. & Moerner, W. E. Rotational Mobility of Single Molecules Affects Localization Accuracy in Super-Resolution Fluorescence Microscopy. Nano Lett 13, 3967–3972 (2013).

41. Engelhardt, J. et al. Molecular Orientation Affects Localization Accuracy in Superresolution Far-Field Fluorescence Microscopy. Nano Lett. 11, 209–213 (2011).

42. Greenspan, P. & Fowler, S. D. Spectrofluorometric studies of the lipid probe, nile red. Journal of Lipid Research 26, 781–789 (1985).

43. Xu, K., Babcock, H. P. & Zhuang, X. Dual-objective STORM reveals three-dimensional filament organization in the actin cytoskeleton. Nat Methods 9, 185–188 (2012).

44. Xu, K., Zhong, G. & Zhuang, X. Actin, Spectrin, and Associated Proteins Form a Periodic Cytoskeletal Structure in Axons. Science 339, 452–456 (2013).

45. Oda, T., Namba, K. & Maéda, Y. Position and Orientation of Phalloidin in F-Actin Determined by X-Ray Fiber Diffraction Analysis. Biophys J 88, 2727–2736 (2005).

46. O’Donnell, L. Mechanisms of spermiogenesis and spermiation and how they are disturbed. Spermatogenesis 4, e979623 (2015).

47. Rathke, C., Baarends, W. M., Awe, S. & Renkawitz-Pohl, R. Chromatin dynamics during spermiogenesis. Biochimica et Biophysica Acta (BBA) - Gene Regulatory Mechanisms 1839, 155–168 (2014).

48. Balhorn, R. The protamine family of sperm nuclear proteins. Genome Biol 8, 227 (2007).

49. Lansac, Y. et al. A route to self-assemble suspended DNA nano-complexes. Sci Rep 6, 21995 (2016).

50. Braun, R. E. Packaging paternal chromosomes with protamine. Nat Genet 28, 10–12 (2001).

51. Orsi, G. A. et al. Biophysical ordering transitions underlie genome 3D re-organization during cricket spermiogenesis. Nat Commun 14, 4187 (2023).

52. Backer, A. S., Lee, M. Y. & Moerner, W. E. Enhanced DNA imaging using super-resolution microscopy and simultaneous single-molecule orientation measurements. Optica, OPTICA 3, 659–666 (2016).

53. Backer, A. S. et al. Single-molecule polarization microscopy of DNA intercalators sheds light on the structure of S-DNA. Science Advances 5, eaav1083 (2019).

54. Zhang, O., Lu, J., Ding, T. & Lew, M. D. Imaging the three-dimensional orientation and rotational mobility of fluorescent emitters using the Tri-spot point spread function. Applied Physics Letters 113, 031103 (2018).

55. Prabhat, P., Ram, S., Ward, E. S. & Ober, R. J. Simultaneous imaging of different focal planes in fluorescence microscopy for the study of cellular dynamics in three dimensions. IEEE Trans Nanobioscience 3, 237–242 (2004).

56. Ram, S., Prabhat, P., Chao, J., Sally Ward, E. & Ober, R. J. High Accuracy 3D Quantum Dot Tracking with Multifocal Plane Microscopy for the Study of Fast Intracellular Dynamics in Live Cells. Biophysical Journal 95, 6025–6043 (2008).

57. Ram, S., Kim, D., Ober, R. J. & Ward, E. S. 3D Single Molecule Tracking with Multifocal Plane Microscopy Reveals Rapid Intercellular Transferrin Transport at Epithelial Cell Barriers. Biophysical Journal 103, 1594–1603 (2012).

58. Abrahamsson, S. et al. MultiFocus Polarization Microscope (MF-PolScope) for 3D polarization imaging of up to 25 focal planes simultaneously. Opt. Express, OE 23, 7734– 7754 (2015).

59. Yan, T., Richardson, C. J., Zhang, M. & Gahlmann, A. Computational correction of spatially variant optical aberrations in 3D single-molecule localization microscopy. Opt. Express, OE 27, 12582–12599 (2019).

60. Sendo, S. et al. Clustering of phosphatase RPTPα promotes Src signaling and the arthritogenic action of synovial fibroblasts. Science Signaling 16, eabn8668 (2023).

61. Blanc, T. et al. Towards Human in the Loop Analysis of Complex Point Clouds: Advanced Visualizations, Quantifications, and Communication Features in Virtual Reality. Front. Bioinform. 1, (2022).

