## Supplementary Information for "Polarization MultiFocus Microscopy for volumetric super-resolution and orientation imaging of biofilaments"

### Supplementary Information Text

#### Supplementary Note 1. Forward and inverse modelling of dipole propagation in PolMFM and retrieval of orientation parameters

This section details the modeling and retrieval of molecular orientation parameters in a PolMFM configuration. We begin with the description of the dipole model, followed by considerations on molecular wobbling. We then derive the mathematical framework for polarized fluorescence intensity and its application in the PolMFM setup and explain the parameter retrieval process.

##### 1. The dipole model

A fluorescent molecule can be described by two distinct transition dipole moments, the absorption dipole $\overrightarrow{\mu_{abs}}$ and the emission dipole $\overrightarrow{\mu_{em}}$. These dipoles are assumed to be aligned[1], and their orientation is described by the unit vector $\vec{\mu}$. In Cartesian coordinates, the orientation of the dipole can be expressed as:

$$\vec{\mu} = \begin{pmatrix} \mu_x \\ \mu_y \\ \mu_z \end{pmatrix} = \begin{pmatrix} \cos(\rho)\sin(\eta) \\ \sin(\rho)\sin(\eta) \\ cos(\eta) \end{pmatrix} \tag{S1}$$

where $\rho$ and $\eta$ denote respectively the in-plane and axial orientation angles (Supplementary Figure S1 A).

The fluorescence intensity recorded on the camera, after propagation through the microscope, is derived based on the model developed in[2]. Assuming isotropic excitation – where all dipoles are excited equally – the formulation considers only the emission signal. Following propagation through an optical system, the amplitude of an electric field propagating along the direction $\vec{k}$, and projected along a given linear polarization direction $\vec{\epsilon}$, can be written at the image plane as:

$$E_{\epsilon}^{img}(\vec{k}) = \left[E_{\epsilon}^{\mu_x}(\vec{k})\ E_{\epsilon}^{\mu_y}(\vec{k})\ E_{\epsilon}^{\mu_z}(\vec{k})\right] \cdot \begin{bmatrix} \mu_x \\ \mu_y \\ \mu_z \end{bmatrix} \tag{S2}$$

where $E_{\epsilon}^{\mu_i}(\vec{k})$ is the component of the electric field propagating along $\vec{k}$, polarized along $\vec{\epsilon}$, and emitted by a dipole oriented along the $i$-axis. The computation of the electric fields $E_{\epsilon}^{\mu_i}(\vec{k})$ will be detailed in Supplementary Note 2 and account for the entire optical system. The intensity at the image plane along the polarization direction $\vec{\epsilon}$ and propagation direction $\vec{k}$ is the square of the electric field and can be derived from eq. (S2) as:

$$I_{\epsilon}(\vec{k}) = \left[E_{\epsilon}^{\mu_x}(\vec{k})^*\ E_{\epsilon}^{\mu_y}(\vec{k})^*\ E_{\epsilon}^{\mu_z}(\vec{k})^*\right] \cdot (\vec{\mu}\vec{\mu}^T) \cdot \begin{bmatrix} E_{\epsilon}^{\mu_x}(\vec{k}) \\ E_{\epsilon}^{\mu_y}(\vec{k}) \\ E_{\epsilon}^{\mu_z}(\vec{k}) \end{bmatrix} \tag{S3}$$

where $\vec{\mu}^T$ is the transpose of $\vec{\mu}$ and $E^*$ is the complex conjugate of $E$.

##### 2. The wobbling and the cone model

In fluorescence experiments, the fluorophore's emission dipole explores multiple orientations due to its rotational flexibility around its linker. This rotational mobility, referred to as "wobbling", happens on timescales of 1-10 ns, much shorter than the integration time of the acquisition camera (~ms)[2,3]. In the model of rotation within a cone[2], the molecule is assumed to visit uniformly each orientation within a cone of solid angle $\delta$ and mean orientation $\{\rho, \eta\}$ (Supplementary Figure S1 B). This dynamic is modelled as the sum of $N$ multiple emission dipoles $\overrightarrow{\mu_n}$, encountered with equal probabilities. The outer product $\left(\overrightarrow{\mu_n}\,\overrightarrow{\mu_n}^T\right)$ of emission dipoles in eq. (S3) can be replaced by a matrix $\boldsymbol{M}$ called the second-order moment matrix:

$$\boldsymbol{M} = \begin{bmatrix} m_{xx} & m_{xy} & m_{xz} \\ m_{xy} & m_{yy} & m_{yz} \\ m_{xz} & m_{yz} & m_{zz} \end{bmatrix} = \frac{1}{N}\sum_{n=1}^{N}\left(\overrightarrow{\mu_n}\,\overrightarrow{\mu_n}^T\right) \tag{S4}$$

It has been shown that $\boldsymbol{M}$ has an eigen-decomposition[2], with two identical eigenvalues noted $\lambda$ and a third distinct eigenvalue, $\lambda'$. As such, the image produced by an emitting dipole rotating within a cone can thus be interpreted as the sum of the images generated by three orthogonal dipoles of amplitudes $|\vec{\mu}|\sqrt{\lambda(\delta)}$ for the two first dipoles, and $|\vec{\mu}|\sqrt{\lambda'(\delta)}$ (Supplementary Figure S1 C). The detailed derivation of the matrix and its eigenvalues is presented in the following subsection.

***Derivation of the second-order moment matrix***

The sum in eq. (S4) can be rewritten as an integration over the cone:

$$\boldsymbol{M} = \frac{|\vec{\mu}|^2}{S}\int_{\phi=0}^{2\pi}\int_{\theta=0}^{\delta/2}\vec{V}\,\vec{V}^{\mathrm{T}}\sin(\theta)\,d\theta d\phi \tag{S5}$$

Here, the normalization factor $\frac{1}{N}$ in eq. (S4) is replaced by the area of the cone solid angle $S = \int_{\phi=0}^{2\pi}\int_{\theta=0}^{\delta/2}\sin(\theta)\,d\theta d\phi = 2\pi(1-\cos\left(\frac{\delta}{2}\right))$. The dipole orientation is parametrized by the angles $\{\phi,\theta\}$ in a rotated coordinate system $\{x',y',z'\}$, where $\theta = 0°$ corresponds to the alignment of the dipole with the mean orientation $\{\rho,\eta\}$ (Supplementary Figure S1 D). The unit dipole moment is rewritten as:

$$\vec{V} = \boldsymbol{R}\begin{bmatrix} \cos(\phi)\sin(\theta) \\ \sin(\phi)\sin(\theta) \\ \cos(\theta) \end{bmatrix} \tag{S6}$$

where a rotation matrix $\boldsymbol{R}(\rho,\eta)$ performs the rotation from the microscope to the cone coordinate system, and is given by:

$$\boldsymbol{R}(\rho,\eta) = \begin{bmatrix} \sin^2(\rho)(1-\cos(\eta)) + \cos(\eta) & \sin(\rho)\cos(\rho)\,(\cos(\eta)-1) & \cos(\rho)\sin(\eta) \\ \sin(\rho)\cos(\rho)\,(\cos(\eta)-1) & \cos^2(\rho)(1-\cos(\eta)) + \cos(\eta) & \sin(\rho)\sin(\eta) \\ -\cos(\rho)\sin(\eta) & -\sin(\rho)\sin(\eta) & \cos(\eta) \end{bmatrix} \tag{S7}$$

Since $\boldsymbol{R}$ is independent of the integration variables $\phi$ and $\theta$, it can be factored out of the integral. The analytical solution of eq. (S5) is:

$$\boldsymbol{M} = |\vec{\mu}|^2\boldsymbol{R}\begin{bmatrix} \lambda(\delta) & 0 & 0 \\ 0 & \lambda(\delta) & 0 \\ 0 & 0 & \lambda'(\delta) \end{bmatrix}\boldsymbol{R}^T \tag{S8}$$

The matrix in the middle is diagonal, and since $\boldsymbol{R}$ is an orthonormal rotation matrix, the eigenvalues of $\boldsymbol{M}$ are simply the diagonal elements of this diagonal matrix. These eigenvalues are explicitly given as[2]:

$$\begin{aligned} \lambda(\delta) &= \frac{1}{6}(1-\cos(\delta/2))(\cos(\delta/2)+2) \\ \lambda'(\delta) &= \frac{1}{3}\frac{(\cos^3(\delta/2)-1)}{(\cos(\delta/2)-1)} \end{aligned} \tag{S9}$$

### 3. Polarized fluorescent intensity

In the previous section, the intensity image produced by a wobbling dipole was derived for a given polarization direction $\vec{\epsilon}$ and a single propagation direction $\vec{k}(\alpha,\beta)$ (eq. (S3)). The intensity at the image plane, $I_{\vec{\epsilon}}$, results from integrating the contributions over all propagation directions $\vec{k}(\alpha,\beta)$ within the acceptance angle of the microscope objective and limited by its numerical aperture. Here, $\alpha \in [0, \alpha_{max} = \sin^{-1}\frac{NA}{n}]$ represents the collection angle, and $\beta \in [0,2\pi]$

corresponds to the in-plane angle (Supplementary Figure S22). After the integration, eq. (S3) becomes:

$$I_{\vec{\epsilon}} = \int_{\beta=0}^{2\pi} d\beta \int_{\alpha=0}^{\alpha_{max}} \left[E_\epsilon^{\mu_x}(\vec{k})^* \; E_\epsilon^{\mu_y}(\vec{k})^* \; E_\epsilon^{\mu_z}(\vec{k})^*\right] \cdot \boldsymbol{M} \cdot \begin{bmatrix} E_\epsilon^{\mu_x}(\vec{k}) \\ E_\epsilon^{\mu_y}(\vec{k}) \\ E_\epsilon^{\mu_z}(\vec{k}) \end{bmatrix} \sin(\alpha)\, d\alpha \tag{S10}$$

Since the second-order moment matrix $\boldsymbol{M}$ depends only on the dipole characteristics $(\rho, \eta, \delta)$, it can be factored out of the integral. This simplifies $I_{\vec{\epsilon}}$ to an inner product:

$$I_{\vec{\epsilon}} = \boldsymbol{K_\epsilon} \cdot \boldsymbol{M} \tag{S11}$$

with $\boldsymbol{K_\epsilon} = [XX_\epsilon \; YY_\epsilon \; ZZ_\epsilon \; XY_\epsilon \; XZ_\epsilon \; YZ_\epsilon]$ and $\boldsymbol{M} = \left[m_{xx} \; m_{yy} \; m_{zz} \; m_{xy} \; m_{xz} \; m_{yz}\right]^T$. The coefficients of $\boldsymbol{K_\epsilon}$ are given by:

$$II_\epsilon = \int_{\beta=0}^{2\pi} d\beta \int_{\alpha=0}^{\alpha_{max}} \left|E_\epsilon^{\mu_i}(\vec{k})\right|^2 \sin(\alpha)\, d\alpha \tag{S12}$$

$$IJ_\epsilon = \int_{\beta=0}^{2\pi} d\beta \int_{\alpha=0}^{\alpha_{max}} 2Re\left\{E_\epsilon^{\mu_i}(\vec{k})^* E_\epsilon^{\mu_j}(\vec{k})\right\} \sin(\alpha)\, d\alpha$$

where $I, J \in \{X, Y, Z\}$.

Using this formalism, the contribution of the single-molecule dipole, represented by $\boldsymbol{M}$, is decoupled from the light propagation through the optical system, represented by $\boldsymbol{K_\epsilon}$.

### 4. Modeling intensity in the PolMFM configuration

The PolMFM configuration splits the signal into six channels corresponding to specific polarization states and focal planes. The output intensities are: $\boldsymbol{I} = [I_{0-}, I_{90-}, I_{45}, I_{135}, I_{0+}, I_{90+}]^T$, where 0° and 90° refer to horizontal and vertical polarizations in the lowest (-) and upper (+) focal planes, and 45° and 135° polarizations correspond to the central focal plane. This can be expressed as a matrix product:

$$\boldsymbol{I} = \begin{bmatrix} I_{0-} \\ I_{90-} \\ I_{45} \\ I_{135} \\ I_{0+} \\ I_{90+} \end{bmatrix} = \begin{bmatrix} XX_{0-} & YY_{0-} & ZZ_{0-} & XY_{0-} & XZ_{0-} & YZ_{0-} \\ XX_{90-} & YY_{90-} & ZZ_{90-} & XY_{90-} & XZ_{90-} & YZ_{90-} \\ XX_{45} & YY_{45} & ZZ_{45} & XY_{45} & XZ_{45} & YZ_{45} \\ XX_{135} & YY_{135} & ZZ_{135} & XY_{135} & ZX_{135} & YZ_{135} \\ XX_{0+} & YY_{0+} & ZZ_{0+} & XY_{0+} & XZ_{0+} & YZ_{0+} \\ XX_{90+} & YY_{90+} & ZZ_{90+} & XY_{90+} & XZ_{90+} & YZ_{90+} \end{bmatrix} \cdot \begin{bmatrix} m_{xx} \\ m_{yy} \\ m_{zz} \\ m_{xy} \\ m_{xz} \\ m_{yz} \end{bmatrix} = \boldsymbol{K} \cdot \boldsymbol{M} \tag{S13}$$

With the use of electric field models (Supplementary Note 2), the coefficients of the propagation matrix $\boldsymbol{K}$ can be computed numerically. The coefficients $XZ_\epsilon$ and $XY_\epsilon$ are found to be equal to zero[4]. It is thus impossible to retrieve the second-order moments $m_{xz}$ and $m_{yz}$. Additionally, all $ZZ_\epsilon$ coefficients are

equal and represent leakage from out-of-plane orientations, with increasing contributions at higher numerical apertures. As shown in[5], the remaining coefficients of $\boldsymbol{K}$ are dependent on only three independent coefficients and the three second-order moments $m_{xx}, m_{yy},$ and $m_{xy}$ can be retrieved. No out-of-plane information can be retrieved, which is consistent with the fact that the setup is performing a projection on the 2D plane of the dipolar information. Thus, eq. (S13) simplifies to:

$$\boldsymbol{I} = \begin{bmatrix} I_{0-} \\ I_{90-} \\ I_{45} \\ I_{135} \\ I_{0+} \\ I_{90+} \end{bmatrix} = \begin{bmatrix} XX_{0-} & YY_{0-} & XY_{0-} \\ XX_{90-} & YY_{90-} & XY_{90-} \\ XX_{45} & YY_{45} & XY_{45} \\ XX_{135} & YY_{135} & XY_{135} \\ XX_{0+} & YY_{0+} & XY_{0+} \\ XX_{90+} & YY_{90+} & XY_{90+} \end{bmatrix} \cdot \begin{bmatrix} m_{xx} \\ m_{yy} \\ m_{xy} \end{bmatrix} = \boldsymbol{K_{2D}} \cdot \boldsymbol{M} \tag{S14}$$

By calibrating $\boldsymbol{K_{2D}}$ experimentally or computing it numerically, the second-order moments ($\boldsymbol{M} = \left[m_{xx} \; m_{yy} \; m_{xy}\right]^T$) can be retrieved using the pseudo-inverse of $\boldsymbol{K_{2D}}$:

$$\boldsymbol{M} = \boldsymbol{K_{2D}^{-1}} \cdot \boldsymbol{I} \tag{S15}$$

### 5. Retrieval of orientation parameters

Using equations (S7), (S8) and (S9), the second-order moments $m_{ij}$ are linked to the dipole's orientation $(\rho, \eta)$ and wobbling $\delta$ by:

$$m_{xx} = |\vec{\mu}|^2 \lambda(\delta)(\cos^2\eta\ \cos^2\rho + \sin^2\rho) + \lambda'(\delta)\ (\sin^2\eta\ cos^2\ \rho) \quad \text{(S16)}$$
$$m_{yy} = |\vec{\mu}|^2 \lambda(\delta)(\cos^2\eta\ \sin^2\rho + \cos^2\rho) + \lambda'(\delta)\ (\sin^2\eta \sin^2\rho)$$
$$m_{xy} = |\vec{\mu}|^2 \sin\rho\cos\rho[\lambda(\delta)(\cos^2\eta - 1) + \lambda'(\delta)\sin^2\eta\ ]$$

Since $\lambda'(\delta) = 1 - 2\lambda(\delta)$, they can be simplified into:

$$m_{xx} = |\vec{\mu}|^2\left[\left(1 - 3\ \lambda(\delta)\right)\ \sin^2\eta\cos^2\rho + \lambda(\delta)\right] \quad \text{(S17)}$$
$$m_{yy} = |\vec{\mu}|^2\left[\left(1 - 3\ \lambda(\delta)\right)\ \sin^2\eta\sin^2\rho + \lambda(\delta)\right]$$
$$m_{xy} = |\vec{\mu}|^2\ \left[\left(1 - 3\ \lambda(\delta)\right)\ \sin^2\eta\ \cos\rho\ \sin\rho\right]$$

We then introduce the three following factors:

$$P_{uv-2D} = \frac{2m_{xy}}{m_{xx}+m_{yy}} = \frac{(1-3\ \lambda)\ \sin^2\eta\ \sin 2\rho}{(1-3\ \lambda)\ \sin^2\eta + 2\ \lambda}, \quad \text{(S18)}$$
$$P_{xy-2D} = \frac{m_{xx}-m_{yy}}{m_{xx}+m_{yy}} = \frac{(1-3\ \lambda)\ \sin^2\eta\ \cos 2\rho}{(1-3\ \lambda)\ \sin^2\eta + 2\ \lambda},$$
$$\lambda_{2D} = \frac{1-P_{2D}}{3-P_{2D}}, \text{ where } P_{2D} = \sqrt{{P_{xy-2D}}^2 + {P_{uv-2D}}^2}.$$

These three factors can be computed with the knowledge of the second-order moments only. In addition, they are related to the mean orientation and wobbling of the emitting dipole $\{\rho, \eta, \delta\}$. However, the simple intensity method analysis developed in this paper is insensitive to out-of-plane orientations as it is equivalent to projecting the dipoles onto a 2D plane. Therefore, it is possible to retrieve the in-plane angle $\rho$ and the 2D projection of the wobbling $\delta_{2D}$ only. Assuming the dipole lies in the sample plane ($\eta = \pi/2$), the orientation and wobbling are determined as:

$$\tan 2\rho = \frac{P_{uv-2D}}{P_{xy-2D}} = \frac{2m_{xy}}{m_{xx} - m_{yy}} \quad \text{(S19)}$$
$$\delta_{2D} = 2\ \mathrm{acos}\left[\frac{-1 + \sqrt{9 - 24\lambda_{2D}}}{2}\right]$$

Thus, the 2D orientation $\rho$ and projected wobbling $\delta_{2D}$ are derived from the second-order moments $\boldsymbol{M}$, obtained by applying the pseudo-inverse of the reduced propagation matrix $\mathbf{K_{2D}}$ to the measured intensities $\boldsymbol{I}$ (eq. (S15)). The orientation parameters $\{\rho, \delta_{2D}\}$ are finally computed from these moments using the analytical expressions of equations (S18) and (S19).

#### Supplementary Note 2. Dipolar emission simulation in PolMFM

The simulation of images generated by wobbling dipoles within both the PolMFM setup or the 4-Polar setup[4] uses the framework outlined in Supplementary Note 1. Intensities in each polarization channel across different focal planes are calculated using eq. (S10). The integration is performed over the entire NA of the objective ($\alpha_{max} = \sin^{-1}(NA/n_1)$ ) and the $\boldsymbol{M}$ matrix is explicitly known derived from equations developed in Supplementary Note 1.2. The only unknown parameters are the electric fields $E_\epsilon^{\mu_i}$, which represent the field components polarized along $\epsilon$ ($\epsilon \in \{x, y\}$) emitted by a fixed dipole oriented along $i = \{x, y, z\}$. These electric fields are computed using a vectorial light propagation model which accounts for the refractive index mismatch at the interface, and that can be found in section 2.3 of [6]. This model propagates the dipolar emission through the interface, collects light via the high-NA objective lens, and determines the amplitude and phase of the light at the BFP of the objective (Supplementary Figure S22).

To simulate the defocus caused by the multifocus grating, the focal plane position is adjusted to three equally spaced planes separated by an interplane distance of $\Delta d = 400\ nm$. This is achieved by applying the following phase shift to the electric fields[7]:

$$\psi_f(\mathrm{d}) = 2\pi \cdot n' \cdot \sqrt{1-(\cos\theta)^2} \cdot \frac{d}{\lambda}$$

where $n' = \begin{cases} n_1 \; if \; d > 0 \text{ (focal plane below the coverslip)} \\ n_2 \; if \; d < 0 \text{ (focal plane above the coverslip)} \end{cases}$ (S20)

To simulate the six channels of the PolMFM, the $x$ and $y$ electric fields are combined as follows to describe the electric fields in the back focal plane produced by a fixed dipole $\overrightarrow{\mu^J}$ :

$\overrightarrow{E^J_{0\pm,BFP}} = \overrightarrow{E^J_x}(\theta,\phi)\; e^{i\left(\psi_{pos}+\psi_f(d_0\pm\Delta z)\right)}$ for the upper (+) and bottom (-) focal plane

$\overrightarrow{E^J_{90\pm,BFP}} = \overrightarrow{E^J_y}(\theta,\phi)\; e^{i(\psi_{pos}+\psi_f(d_0\pm\Delta z))}$ for the upper (+) and bottom (-) focal plane

$\overrightarrow{E^J_{45,BFP}} = \frac{1}{\sqrt{2}}\left(\overrightarrow{E^J_x}(\theta,\phi)\; e^{i\left(\psi_{pos}+\psi_f(d_0)\right)} + \overrightarrow{E^J_y}(\theta,\phi)\; e^{i\left(\psi_{pos}+\psi_f(d_0)\right)}\right)$ for the central plane (S21)

$\overrightarrow{E^J_{135,BFP}} = \frac{1}{\sqrt{2}}\left(-\overrightarrow{E^J_x}(\theta,\phi)\; e^{i\left(\psi_{pos}+\psi_f(d_0)\right)} + \overrightarrow{E^J_y}(\theta,\phi)\; e^{i\left(\psi_{pos}+\psi_f(d_0)\right)}\right)$ for the central plane

Here $d_0$ is the position of the central focal plane and $\psi_{pos}$ is the phase shift to account for the emitter lateral and axial position, as specified in [7].

To simulate the four polarization channels of the 4-Polar scheme[4], eq. (S21) were also used with the only difference that no defocus was applied ($\Delta z = 0$) since all the 4 polarization channels lye at the same focal plane.

The electric fields in the BFP are Fourier transformed using the Fast Fourier Transform (FFT) algorithm in Matlab to account for the propagation through the tube lens of the microscope and provide the fields at the camera for each polarization channel and focal planes. According to the $\boldsymbol{M}$ matrix model for wobbly dipole developed in Supplementary Note 1.2, the image of a molecule rotating within a cone is identical to an image of three fixed orthogonal super-imposed dipoles $\{\overrightarrow{\mu^1}, \overrightarrow{\mu^3}, \overrightarrow{\mu^3}\}$ , of amplitudes $\mu_1 = |\vec{\mu}|\sqrt{\lambda'(\delta)}$ and $\mu_2 = \mu_3 = |\vec{\mu}|\sqrt{\lambda(\delta)}$, given by eq. (S9). Therefore, the simulated intensities are finally the sum of the squared electric fields of each dipole component:

$$I_{0\pm} = \sum_{j=1}^{3} |FFT\{\overrightarrow{E^J_{0\pm,BFP}}\}|^2; I_{90\pm} = \sum_{j=1}^{3} |FFT\{\overrightarrow{E^J_{90\pm,BFP}}\}|^2 \qquad \text{(S22)}$$

with (+) representing the upper focal planes and (-) the bottom ones,

and

$$I_{45} = \sum_{j=1}^{3} |FFT\{\overrightarrow{E^J_{45,BFP}}\}|^2; I_{135} = \sum_{j=1}^{3} |FFT\{\overrightarrow{E^J_{135,BFP}}\}|^2$$

for the two central focal planes.

Since the tube lens of the microscope performs a Fourier transform operation[8], this final operation (eq. (S22)) corresponds to the propagation of the electric field through the tube lens. The parameters of the PolMFM microscope that were plugged into the simulations can be found in Table S1.

To correctly match the pixel size of the simulated PSFs with our optical system, we must account for the scaling introduced by both the tube lens and the relay lens system. The tube lens performs the Fourier transform with a scaling factor $\frac{1}{\lambda f_{TL}}$, while the relay lens system, which conjugates the primary image plane with the camera, introduces an additional scaling factor $M = f_2/f_1$, where $f_1$ and $f_2$ are the focal lengths of the relay lenses[8,9]. Initially, the electric fields (eq. (S21)) are sampled on a cartesian grid $(X, Y)$ representing the BFP, within a range $x \in [x_{min}, x_{max}]$ and $y \in [y_{min}, y_{max}]$,using a regular sampling interval of $\Delta x$ and $\Delta y$. To properly account for the circular aperture of the objective's BFP, the electric fields are set to zero outside the numerical aperture, defined by $\sqrt{X^2+Y^2} > f \cdot \sin\alpha_{max}$, where $f = f_{TL}/OM$ is the focal length of the objective and $\alpha_{max} = \mathrm{asin}(NA/n_1)$. When applying the FFT algorithm, the field of view (FOV) in the image plane is determined by the inverse of the pixel size in the BFP, while the pixel size in the image plane corresponds to the inverse of the FOV in the BFP. Consequently, to ensure correct image scaling and pixel size, the BFP grid must be extended with zero-padding before performing the FFT. The required padding size, defined as the number of additional samples on each side of the grid, is given by $N_{pad} = \lfloor \frac{\lambda \cdot f_{TL} \cdot M - px\ size_{ccd} \cdot (x_{max} - x_{min})}{2 \cdot \Delta x \cdot px\ size_{ccd}} \rfloor$, ensuring that both the tube lens $f_{Tl}$ scaling and the relay lenses $M$ magnification are correctly incorporated.

**Supplementary Note 3. MFM design and performance**

PolMFM is based on the implementation of a three-plane multifocus microscopy (MFM) configuration in addition to the polarization projection. MFM operates by inserting diffractive optical elements into the emission path of a widefield fluorescence microscope, enabling the simultaneous acquisition of multiple focal planes. Central to this approach is the multifocus grating (MFG), which separates the emission light into a 2D array of diffraction orders.

For PolMFM, the MFG's central motif was specifically optimized to achieve maximum diffraction efficiency and uniformity in the three-by-one central diffraction orders. A binary phase grating design was chosen for ease of fabrication. The optimization procedure described in [10] was applied to ensure multicolor compatibility across the visible spectrum (530–670 nm), while maximizing diffraction uniformity and efficiency.

Figure S6-A illustrates the grating motif, where white and black regions correspond to different etching depths in the fused silica substrate. These depths produce a relative phase shift of $2.15\ \pi/2$ between light paths, corresponding to an etching depth of 625 nm when accounting for the refractive index of fused silica. Theoretical performance simulations predicted a diffraction efficiency of 67% and a uniformity of 96% Figure S6-B. In practice, the fabricated grating achieved 64% efficiency and 92% uniformity.

Each motif of the grating was designed to span an area of 12µm (the grating period) and was tiled to fully cover an equivalent size of the back focal plane (BFP) of the objective lens, considering the demagnification introduced by intermediate optics. To generate defocus between the diffraction orders, a radial distortion was introduced in the grating periodicity Supplementary Figure S6-C, as described in [10–12]. This defocus corresponds to an interplane separation of 370 nm at an imaging wavelength of 600 nm. Figure S6 presents the grating design, along with measured performance parameters.

**Supplementary Note 4. Calibrations: channel registration, intensity calibration and polarization calibration**

The raw data acquired with the PolMFM consists of a 6 channels image, each channel arising from a different polarization and focal plane combination. To reconstruct a 3D polarized stack from the raw data, the 6 channels need to be carefully aligned, using a registration matrix (Supplementary Note 4.3). Moreover, as the PolMFM does not split fluorescence with equal intensity, the reconstruction of the 6-channels stack is corrected with the help of correction factors that are pre-calibrated (Supplementary Note 4.2). Finally, the propagation matrix $\boldsymbol{K_{2D}}$ must be experimentally built to ensure that it considers experimental deviations from perfect setup, such as the imbalance of intensity transmitted in the different focal planes or polarization distortions (Supplementary Note 4.3).

1. **Channel registration**

A piezoelectric stage displaces axially a 100 nm fluorescent beads sample in 100 nm increments to acquire a focal series using the PolMFM. During the scan, the same beads appear successively in focus on each channel. To precisely localize the position of all the beads in each channel, a maximum intensity projection of the z-stack is computed (Supplementary Figure S3-A), and the beads are localized by fitting the intensity in each channel to a 2D Gaussian profile (circles in Supplementary Figure S3-A). The super-localizations are assigned to their corresponding channels (colors Supplementary Figure S3-A), and a Procrustes algorithm (provided in MATLAB) is used to find the optimal transformation matrix that minimizes the sum of squared differences between the super-localizations of each channel. The 6 channels of each raw image are aligned on top of each other using the transformation matrix to build a 6-channel stack (Fig. 1-C-2).

The z-stack of fluorescent beads is also used to compute the interplane distance of the MFM. The axial intensity profile of the beads in each channel of the PolMFM is extracted from the z-stack (colored lines in Supplementary Figure S3-B). For each channel, the axial position of the best focus is computed by a Gaussian fitting of the intensity profiles (gray dashed lines in Supplementary Figure S3-B). The axial position of each focal plane of the MFM is taken as the mean fitted z-position of the two perpendicular polarizations. The retrieved z-positions of the three planes are fitted with a linear curve, and the slope directly gives the interplane distance (Supplementary Figure S3-D). This interplane distance is later used as the axial pixel size for the 3D Gaussian fitting on real single molecule data.

#### 2. Intensity correction factors

The different optical elements on the emission path will introduce some intensity difference between the 6 recorded channels. As our method relies on the fluorescence intensity measurement to retrieve the required information, such intensity differences should be considered. To calibrate the intensity imbalance between the different channels, we acquire an image using the unpolarized white lamp of the microscope (Supplementary Figure S4**Error! Reference source not found.**). The intensity calibration factors $C_i$ ($i \in \{0-, 90-, 45, 135, 0+, 90+\}$) are built by normalizing the measured intensity in each channel to the intensity of the brightest channel. At a later step, during the 6-channel stack reconstruction stage, each channel image (Fig. 1-C-2) is normalized by its correction factor $C_i$.

#### 3. Propagation matrix calibration

The propagation matrix $\boldsymbol{K_{2D}}$ matrix is built by consecutively cancelling some second-order coefficients $m_{ij}$ and acquiring dipolar emissions of known orientations. For a fixed dipole ($\delta = 0°$) lying in the sample plane ($\eta = 90°$), the second order dipole moments are, according to eq (S9) and (S17):

$$\begin{aligned} m_{xx}(\rho, \eta = 90°, \delta = 0°) &= |\vec{\mu}|^2 \cos^2 \rho \\ m_{yy}(\rho, \eta = 90°, \delta = 0°) &= |\vec{\mu}|^2 \sin^2 \rho \\ m_{xy}(\rho, \eta = 90°, \delta = 0°) &= |\vec{\mu}|^2 \, cos\, \rho \sin \rho \end{aligned} \qquad \text{(S23)}$$

As such, each coefficient $m_{ij}$ can be isolated, allowing the measured intensities $\boldsymbol{I}$ to be related to a specific column of $\boldsymbol{K_{2D}}$. To mimic dipoles oriented in the sample plane, we used the unpolarized transmission lamp of the microscope (LED) to illuminate the sample plane at which a linear polarizer is placed, and we collected the light filtered at the appropriate wavelength on the camera. The polarizer is placed in a rotative mount to control the orientation $\rho$ of the mimicked in-plane dipole. The images obtained are shown in Figure S5 for a polarizer oriented along 0° (top), 90° (middle) and 45° (bottom). According to equation (S23), the only second-order dipole moment that does not vanish for $\rho = 0°$ is $m_{xx}$, allowing to compute the first column of the matrix $\boldsymbol{K_{2D}}$ by normalizing the intensities measured in each channel with the normalization factor $|\vec{\mu}|^2 = \sum_j I_j$ (Supplementary Figure S5, top row). In a similar way, the second column of $\boldsymbol{K_{2D}}$ is retrieved by rotating the polarizer to $\rho = 90°$ such as the only non-vanishing coefficient is $m_{yy}$ (Supplementary Figure S5, middle row). Finally, the polarizer is rotated to $\rho = 45°$ to equalize the three coefficients $m_{xx} = m_{yy} = m_{xy} = |\vec{\mu}|^2/2$ and compute the third column of $\boldsymbol{K_{2D}}$ using the previously retrieved two first columns (Supplementary Figure S5, bottom row).

### Supplementary Note 5. Quantifications performed on silica beads samples.

For each PolMFM acquisition of a silica bead sample, a widefield z-stack of the corresponding field of view (FOV) was pre-acquired before single molecule imaging. The MATLAB function ‘imfindcircles’ (a Circular Hough Transform (CHT) based algorithm for finding circles in images) was used on a maximum intensity projection of the widefield z-stack to identify the different beads (Supplementary Figure S13-A). The central slice of the bead was inferred by computing Tamura’s coefficients[13] at each slice, then fitting a Gaussian curve on the resulting coefficients. The position of the Gaussian curve's maximum was considered as the central z slice of the bead (Supplementary Figure S13-B). The bead radius was then determined using ‘imfindcircles’ on the central slice image.

Next, the lateral bead center $(x_0, y_0)$ was computed directly on the point cloud of the SMOLM acquisition. First, the center of mass of the beads is taken as the initial guess for the center. The number of detections within concentric annulus rings of 300 nm width was computed (Supplementary Figure S13-C, top) and plotted as a function of increasing radius (Supplementary Figure S13-C, bottom). The ‘findpeaks’ function in Matlab was used to calculate the peak width of this function. A nonlinear minimization (Matlab function ‘fminsearch’) was applied to minimize the width of the distribution while varying the center $(x_0, y_0)$.

Using the determined bead center position, and based on the expected radiality of molecular orientation, the theoretical orientation angles $\rho_{theo}$ were calculated for each detected molecule from its 2D coordinates relative to the bead center $(x_0, y_0)$:

$$\rho_{theo} = \tan^{-1}\left(\frac{y - y_0}{x - x_0}\right)$$

To allow for comparison and pooling of data across different beads, the 2D coordinates were then normalized by the bead radius, ensuring that all beads had a normalized radius equal to one. This normalization enabled concatenation of results from multiple beads for further analysis:

$$x' = \frac{x - x_0}{radius}, y' = \frac{y - y_0}{radius}, z' = \frac{z}{radius}$$

### Supplementary Figures

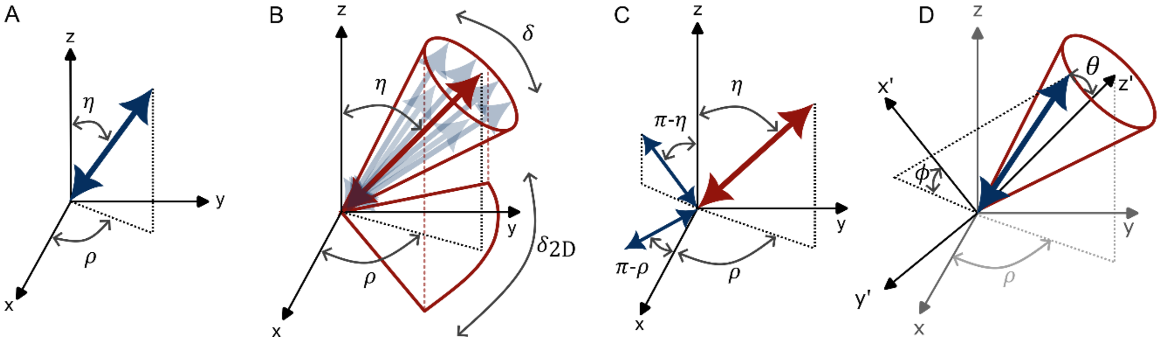


**Supplementary Figure S1. Dipole parametrization and modelling.** A) Dipole angular parametrization with azimuthal angle $\boldsymbol{\rho}$ and polar angle $\boldsymbol{\eta}$. Z represents the optical axis. B) A rotating dipole is parametrized by its mean orientation $\{\boldsymbol{\rho}, \boldsymbol{\eta}\}$ and wobbling within a cone of aperture angle $\boldsymbol{\delta}$. C) Parametrization of the wobbling within a cone model with the help of three fixed orthogonal dipoles of amplitude $|\vec{\boldsymbol{\mu}}|\sqrt{\boldsymbol{\lambda}'(\boldsymbol{\delta})}$ (in red) and $|\vec{\boldsymbol{\mu}}|\sqrt{\boldsymbol{\lambda}(\boldsymbol{\delta})}$ (in blue). D) Rotated coordinate system $\{\boldsymbol{x}', \boldsymbol{y}', \boldsymbol{z}\}$ where the mean orientation of the dipole is parametrized with azimuthal angle $\boldsymbol{\phi}$ and polar angle $\boldsymbol{\theta}$.

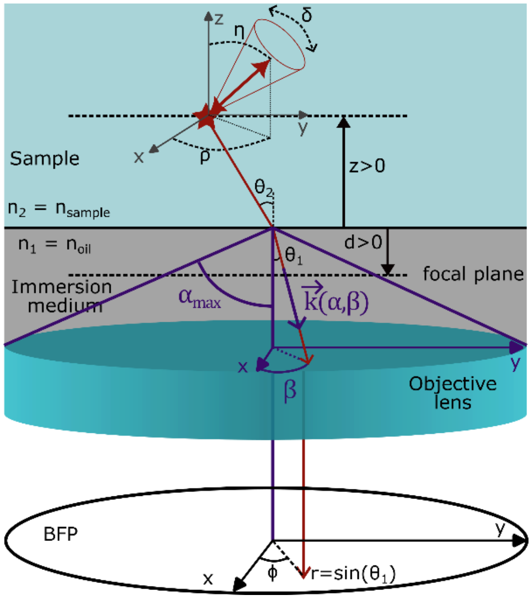


**Supplementary Figure S2. Parametrization of the propagation in the objective lens and back focal plane.** Inspired from [4,7].

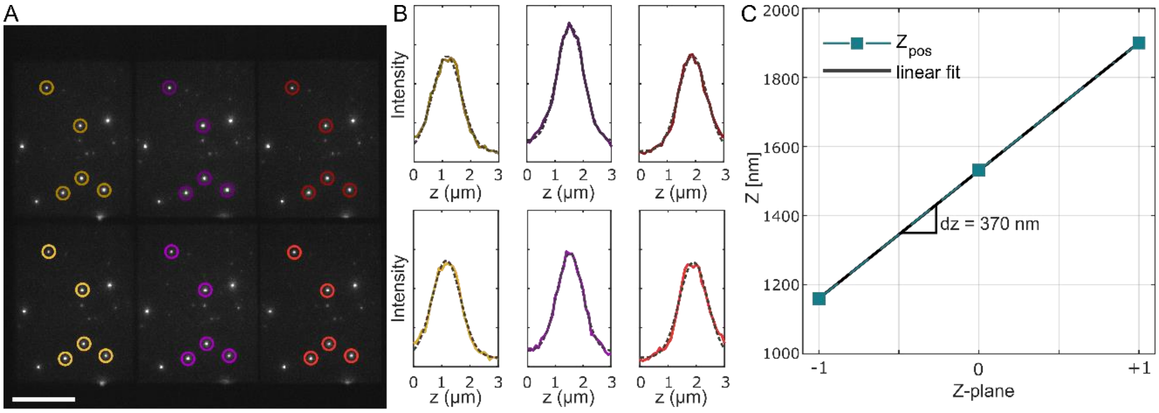


**Supplementary Figure S3. Image channels registration and Z-plane calibration.** A) Maximum intensity projection of a z-stack of the fluorescence signal of 200 nm diameter beads displaced axially with steps of 100 nm and excited with a 560 nm laser. The circles represent the localizations of the beads in each channel that are used for generating the different channel registration matrices. Scale bar = 10 µm. B) Axial intensity profiles of one bead in the 6 different channels (colored line) and corresponding fit (dashed grey line) for determining the z-position of the best focus of each channel, taken as the center of the fitted Gaussian.CD) Plot of the z-position of each focal plane (squares), taken as the means of the best positions retrieved in the two perpendicular polarized channels. Linear fit (black line) to infer the interplane distance **ΔZ** between each focal plane of the PolMFM.

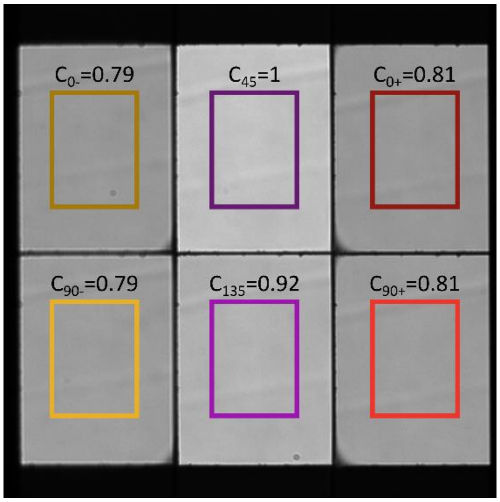


**Supplementary Figure S4. White-lamp acquisition for intensity correction factor calibration with 617±73 nm emission filter.** $\boldsymbol{C_i}$ are the correction factors, computed as the mean intensity $\boldsymbol{I_i}$ in the rectangle area of each channel, were $\boldsymbol{i} \in \{\mathbf{0-}, \mathbf{90-}, \mathbf{45}, \mathbf{135}, \mathbf{0+}, \mathbf{90+}\}$ $i\in\{0-,90-,45,135,90+,135+\}$, normalized by the maximum intensity $\boldsymbol{I_{max}}$.

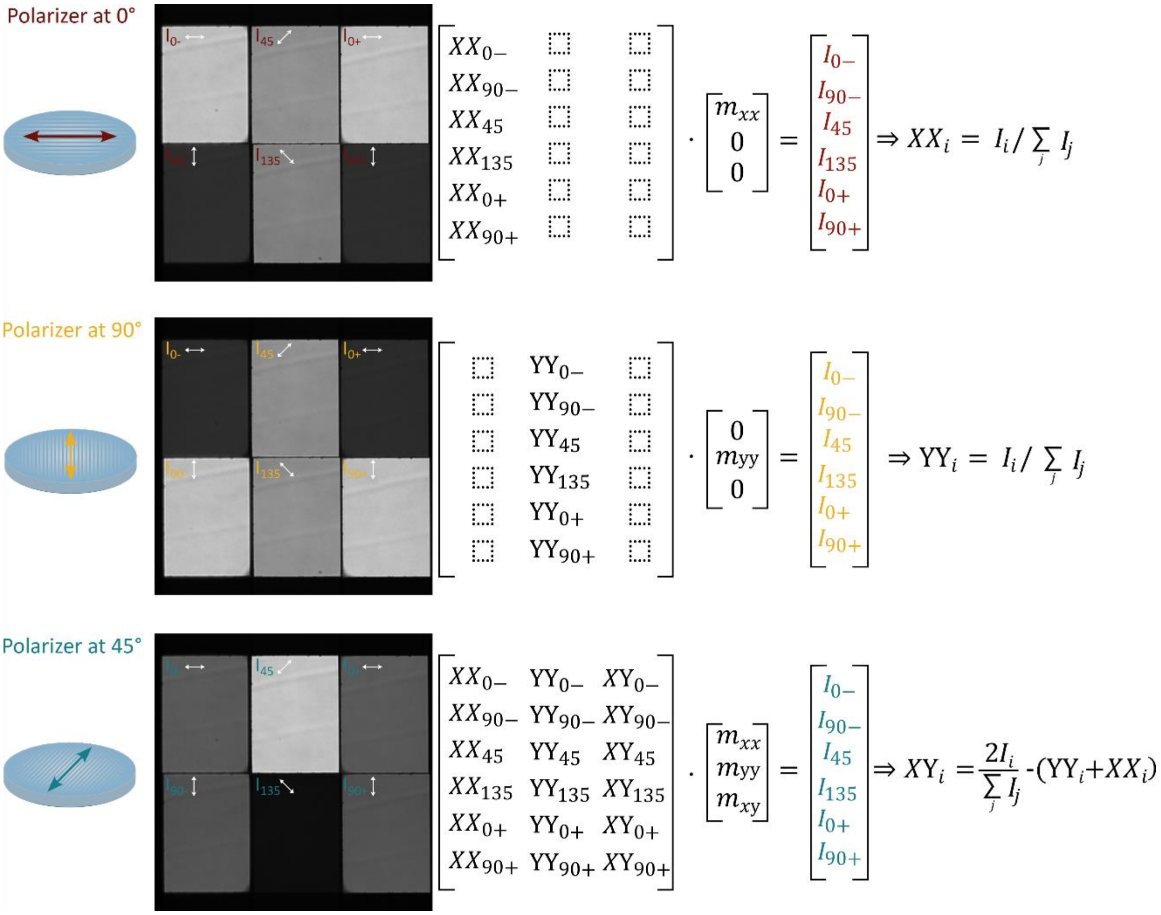


**Supplementary Figure S5. Procedure for $K$ matrix calibration.** The 3 images correspond to an acquisition in the PolMFM with the microscope LED illumination, filtered for the emission wavelength at 617±76 $nm$, and polarized at the sample plane with a linear polarizer. The polarizer orientation varies: 0° (top), 90° (middle) and 45° (bottom). The equations (right) recall the procedure for the computation of the coefficients of the propagation matrix.

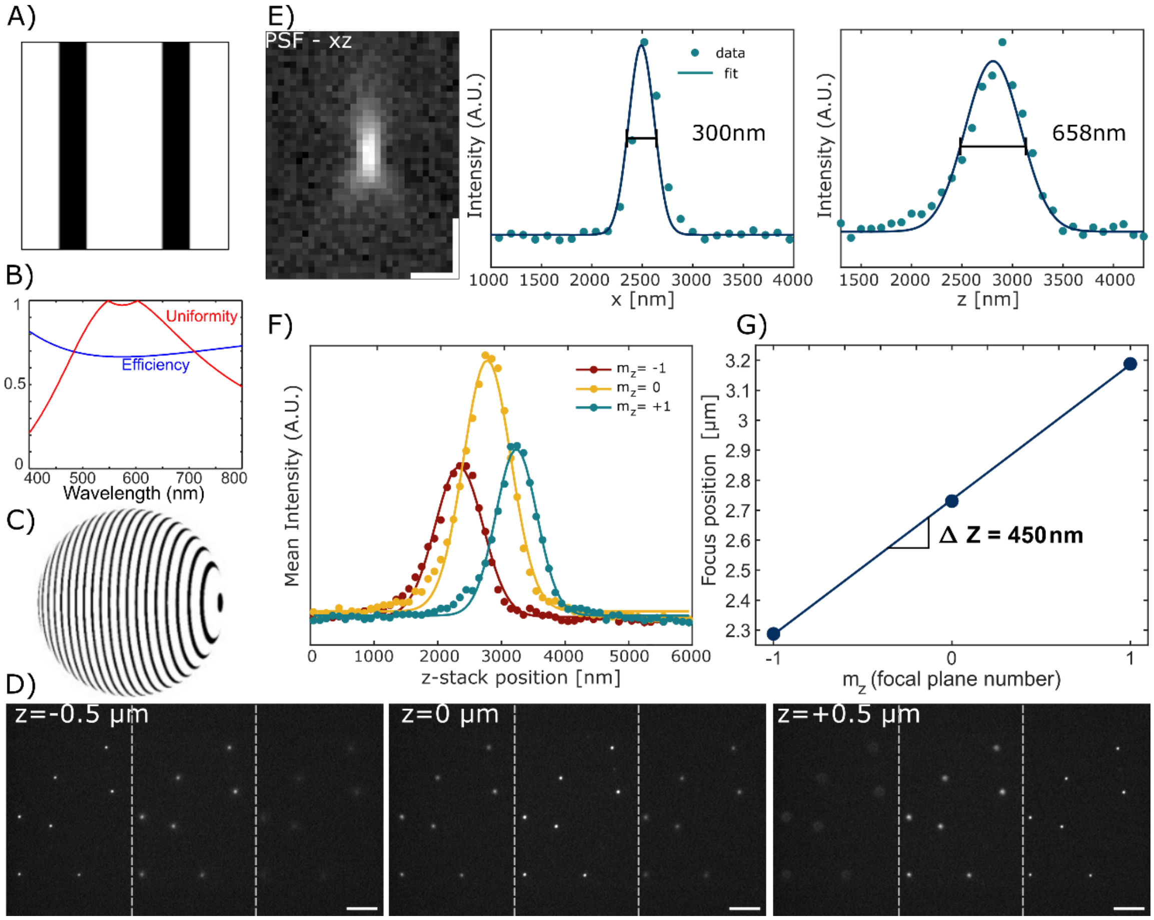


**Supplementary Figure S6. Design and characterization of the multifocal grating.** A) Grating motif as generated by simulation resulting in a 3-by-1 diffraction pattern. B) Theoretical dependency of the diffraction uniformity and efficiency of the designed grating motif as function of the imaging wavelength. C) exaggerated view of the MFG design showing the main motif allowing the 3x1 diffraction of the emission with high efficiency and uniformity. The distortion in the motif is required to introduce different defocusing powers in each of the diffraction orders. D) fluorescent beads deposited on a coverglass are scanned along the optical axis to characterize the MFM microscope performances. Scale bar = 5 µm. E) Characterization of the 3D PSF shape and size showing diffraction-limited extent as expected. Scale bar = 1 µm. F) The axial displacement of the bead allows to compute the intensity profile of the beads for each plane as function of the displacement. G) The peak position of the fluorescence intensity for each MFM plane is plotted as function of Z to retrieve the exact interplane spacing of the MFM.

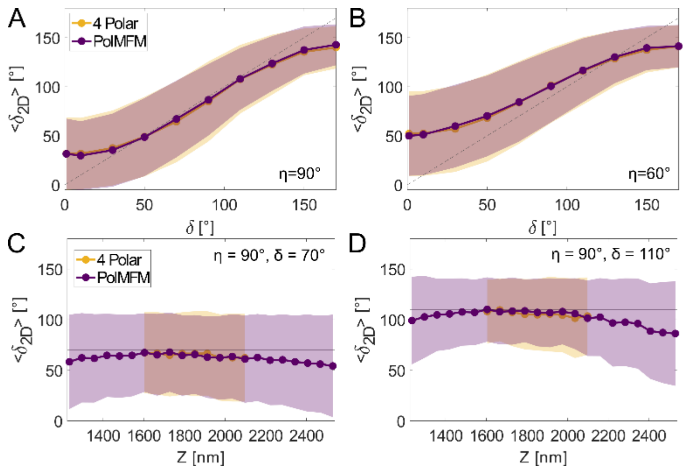


**Supplementary Figure S7. Bias and error on $\boldsymbol{\delta_{2D}}$ estimation.** For each simulated dipole of given orientation $\{\rho, \eta, \delta\}$, 100 independent iterations of noise are generated. Each molecule emits 1000 photons and a background of 2 photons per pixel is added to the images. The graphs show the mean (dots) ± standard deviations (shaded areas) of the parameters retrieved over the 100 iterations, in the PolMFM case (purple) and in-plane 4-Polar setup (yellow). A) Retrieved $\boldsymbol{\delta_{2D}}$ as a function of wobbling $\boldsymbol{\delta}$ for in-plane dipole ($\boldsymbol{\eta} = \mathbf{90}°$) located axially at the central focal plane. Results are averaged for varying azimuthal orientation ($\boldsymbol{\rho} = \mathbf{0}°: \mathbf{15}°: \mathbf{165}°$). B) Same as A for inclined dipoles ($\boldsymbol{\eta} = \mathbf{60}°$). C) Retrieved $\boldsymbol{\delta_{2D}}$ as a function of axial position $\boldsymbol{Z}$ for in-plane dipole ($\boldsymbol{\eta} = \mathbf{90}°$) with wobbling $\boldsymbol{\delta} = \mathbf{70}°$. Results are averaged for varying azimuthal orientation ($\boldsymbol{\rho} = \mathbf{0}°: \mathbf{15}°: \mathbf{165}°$). D) Same as C with wobbling $\boldsymbol{\delta} = \mathbf{110}°$.

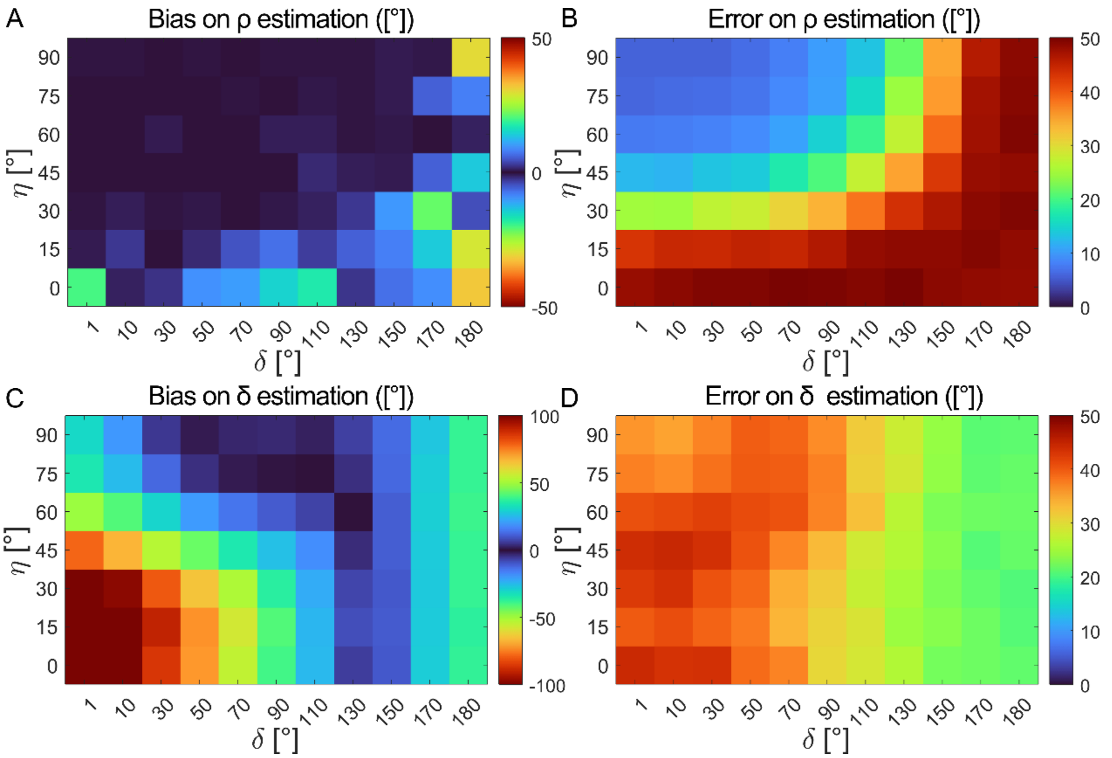


**Supplementary Figure S8. Bias and error matrices on orientation parameter retrieval as a function of wobbling $\boldsymbol{\delta}$ and polar angle $\boldsymbol{\eta}$.** Each pixel represents parameter retrieval for a dipole located at the central focal plane with given polar angle $\boldsymbol{\eta}$, given wobbling $\boldsymbol{\delta}$ and azimuthal orientation varied within $\boldsymbol{\rho = 0: 15: 165°}$. For each simulated dipole of given orientation $\{\rho, \eta, \delta\}$, 100 independent iterations of noise are generated. Each molecule emits 1000 photons and a background of 2 photons per pixel is added to the images. A) Bias on $\boldsymbol{\rho}$ estimation. B) Error on $\boldsymbol{\rho}$ estimation. C) Bias on $\boldsymbol{\delta_{2D}}$ estimation. D) Error on $\boldsymbol{\delta_{2D}}$ estimation.

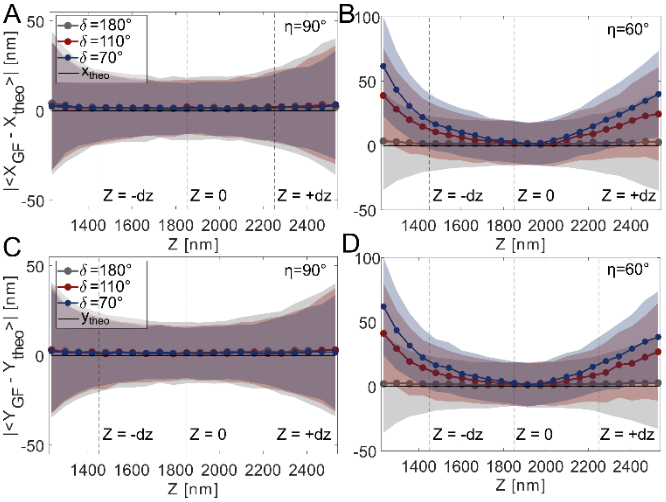


**Supplementary Figure S9. Bias and error on lateral localization as a function of axial position.** For each simulated dipole of given orientation $\{\rho,\ \eta,\ \delta\}$, 100 independent iterations of noise are generated. Each molecule emits 1000 photons and a background of 2 photons per pixel is added to the images. The graphs show the mean (dots) ± standard deviations (shaded areas) of the parameters retrieved over 100 iterations. A) Absolute bias on $\boldsymbol{x} \pm$ standard deviation as a function of axial position of the dipole $\boldsymbol{Z}$ for in-plane dipoles ($\boldsymbol{\eta} = \mathbf{90}°$), with wobbling $\boldsymbol{\delta} = \mathbf{180}°$ (gray), $\boldsymbol{\delta} = \mathbf{110}°$ (red) or $\boldsymbol{\delta} = \mathbf{70}°$ (blue). Results are averaged for varying azimuthal orientation ($\boldsymbol{\rho} = \mathbf{0}°\colon \mathbf{15}°\colon \mathbf{165}°$). B) Same as A for inclined dipoles ($\boldsymbol{\eta} = \mathbf{60}°$). C) Same as A on $\boldsymbol{y}$ retrieval. D) Same as B on $\boldsymbol{y}$ retrieval.

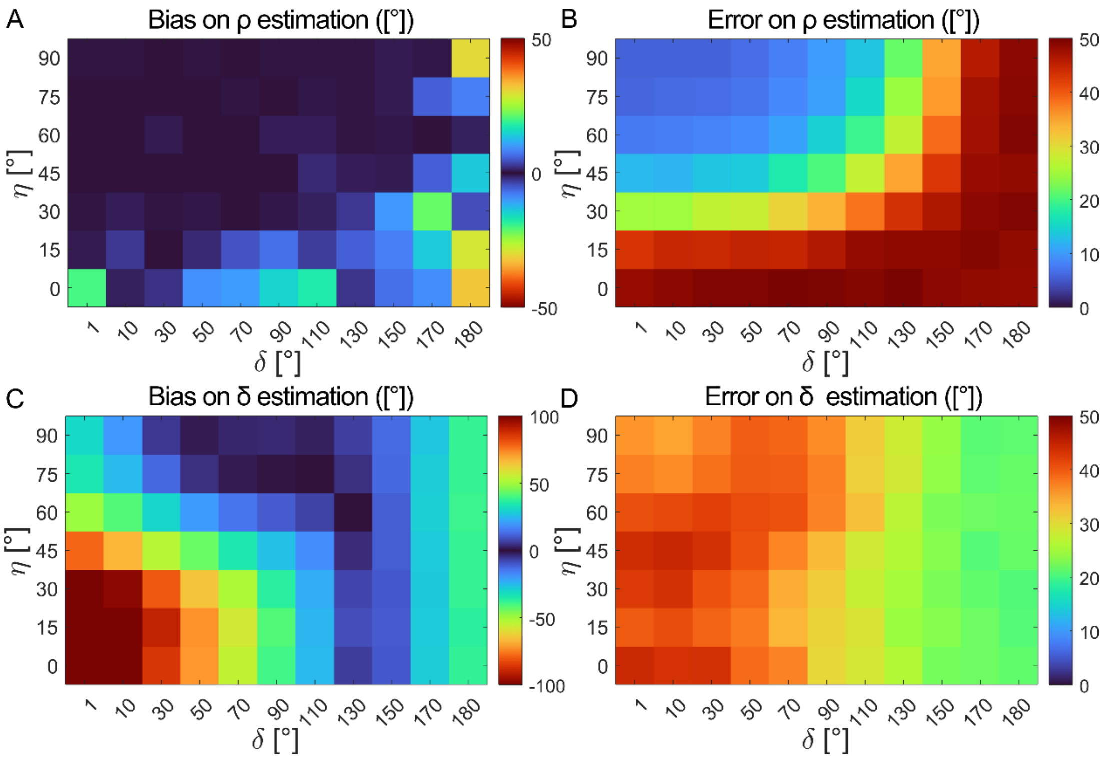


**Supplementary Figure S10. Bias and error maps on lateral localization as a function of wobbling $\boldsymbol{\delta}$ and polar angle $\boldsymbol{\eta}$.** Each pixel represents parameter retrieval for a dipole located at the central focal plane with given polar angle $\boldsymbol{\eta}$, given wobbling $\boldsymbol{\delta}$ and azimuthal orientation varied within $\boldsymbol{\rho = 0{:}\,15{:}\,165°}$. For each simulated dipole of given orientation $\{\rho,\ \eta,\ \delta\}$, 100 independent iterations of noise are generated. Each molecule emits 1000 photons and a background of 2 photons per pixel is added to the images. A) Bias on $\boldsymbol{x}$ estimation. B) Bias on $\boldsymbol{y}$ estimation. C) Error on $\boldsymbol{x}$ estimation. D) Error on $\boldsymbol{y}$ estimation.

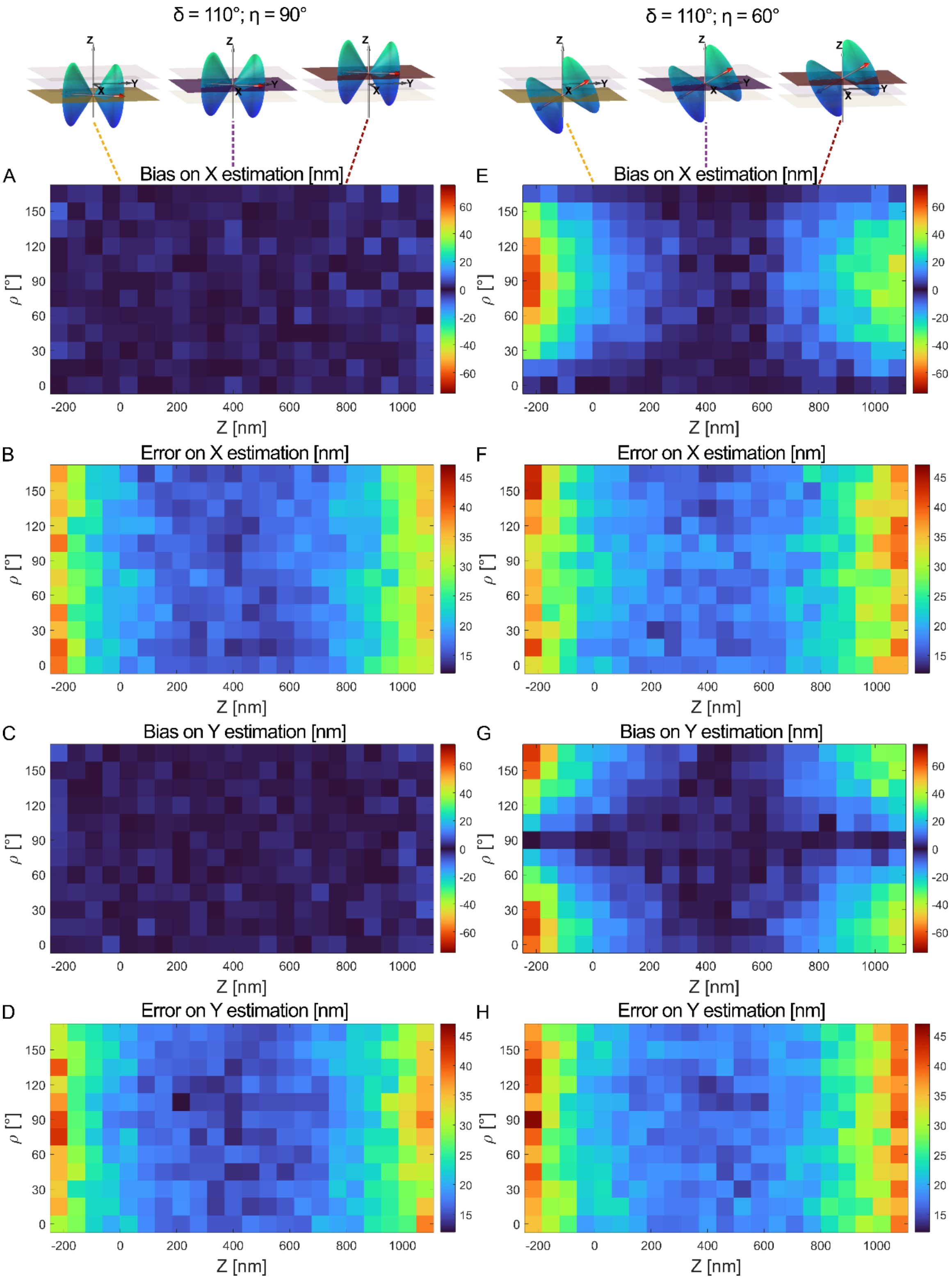


**Supplementary Figure S11. Bias and error maps on lateral localization as a function of axial position and azimuthal orientation.** Each pixel represents parameter retrieval for a dipole with a given polar angle $\boldsymbol{\eta}$ and wobbling $\boldsymbol{\delta}$. For each simulated dipole of given orientation $\{\rho, \eta, \delta\}$ and $\boldsymbol{Z}$ position, 100 independent iterations of noise are generated. Each molecule emits 1000 photons and a background of 2 photons per pixel is added to the images. A) Bias on $\boldsymbol{x}$ for $\boldsymbol{\delta} = \mathbf{110}°$ and $\boldsymbol{\eta} = \mathbf{90}°$. B) Error on $\boldsymbol{x}$ for $\boldsymbol{\delta} = \mathbf{110}°$ and $\boldsymbol{\eta} = \mathbf{90}°$.

C) Bias on $\boldsymbol{y}$ for $\boldsymbol{\delta = 110}°$ and $\boldsymbol{\eta = 90}°$. D) Error on $\boldsymbol{y}$ for $\boldsymbol{\delta = 110}°$ and $\boldsymbol{\eta = 90}°$**.** E) Bias on $\boldsymbol{x}$ for $\boldsymbol{\delta = 110}°$ and $\boldsymbol{\eta = 60}°$. F) Error on $\boldsymbol{x}$ for $\boldsymbol{\delta = 110}°$ and $\boldsymbol{\eta = 60}°$. G) Bias on $\boldsymbol{y}$ for $\boldsymbol{\delta = 110}°$ and $\boldsymbol{\eta = 60}°$. H) Error on $\boldsymbol{y}$ for $\boldsymbol{\delta = 110}°$ and $\boldsymbol{\eta = 60}°$**.**

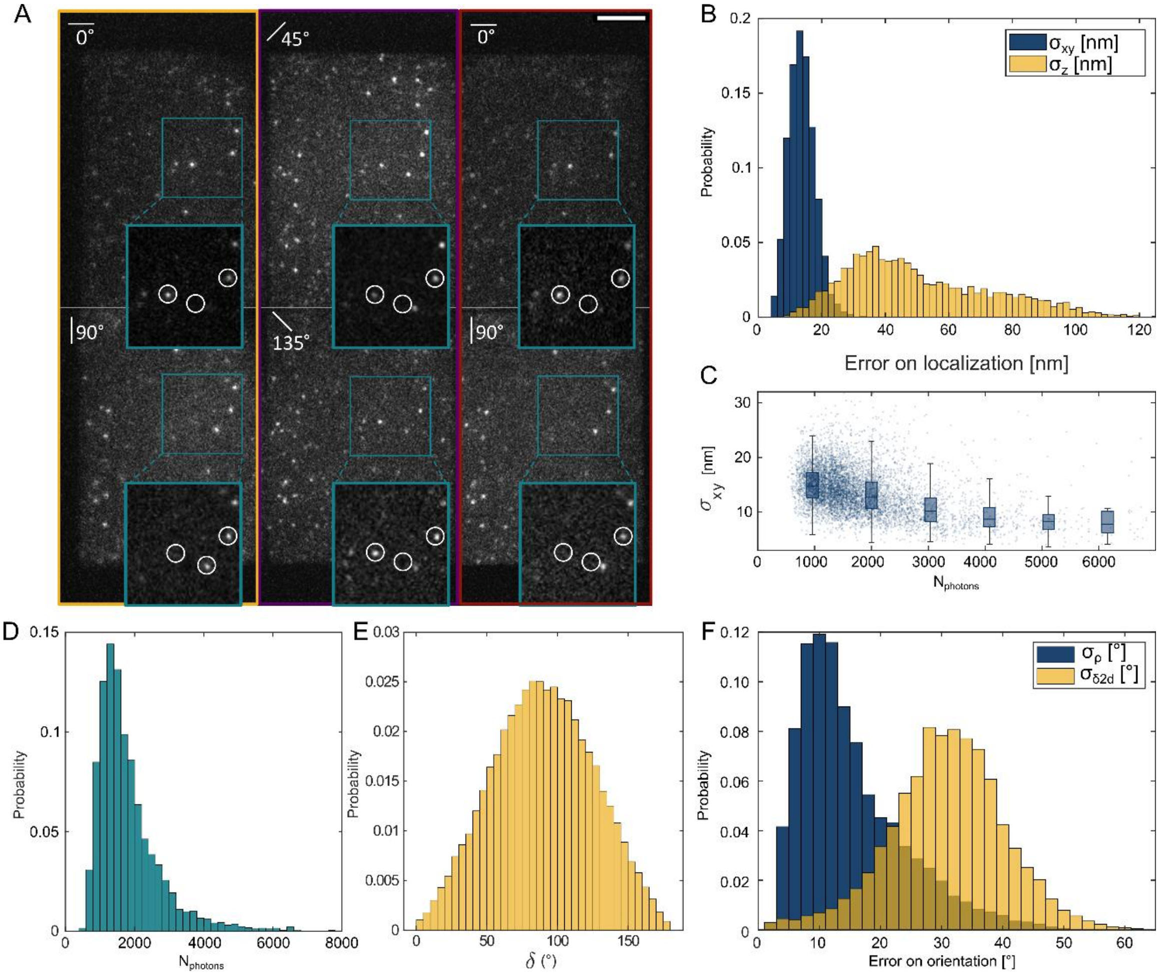


**Supplementary Figure S12. Quantifications on PolMFM imaging of JF 549 dyes embedded in PVA.** A) Raw data. Blue squares: zoom on a few molecules. The 3 circled molecules appear brightly on different polarization channels, indicating different orientations. B) Histograms of error on $\mathbf{x}$ and $\mathbf{y}$ localizations. The error is taken as the standard deviation of the retrieved value over multiple detections of the same molecule, where molecules appearing between 10 and 30 times are kept. C) Error on lateral localization as a function of number of detected photons. In the scatter plots, each point represents a molecule. In the box plots the statistics is computed for molecules withing a range of photon numbers. D) Histogram of mean photon number of all detected molecules, for which the mean value is taken on all the detections originating from the same molecule. E) Histogram of retrieved wobbling $\boldsymbol{\delta_{2D}}$ for all the molecules. F) Histograms of error on $\boldsymbol{\rho}$ and $\boldsymbol{\delta_{2D}}$ estimation. The error is taken as the standard deviation of the retrieved value over multiple detections of the same molecule, where molecules appearing at least 10 times are kept.

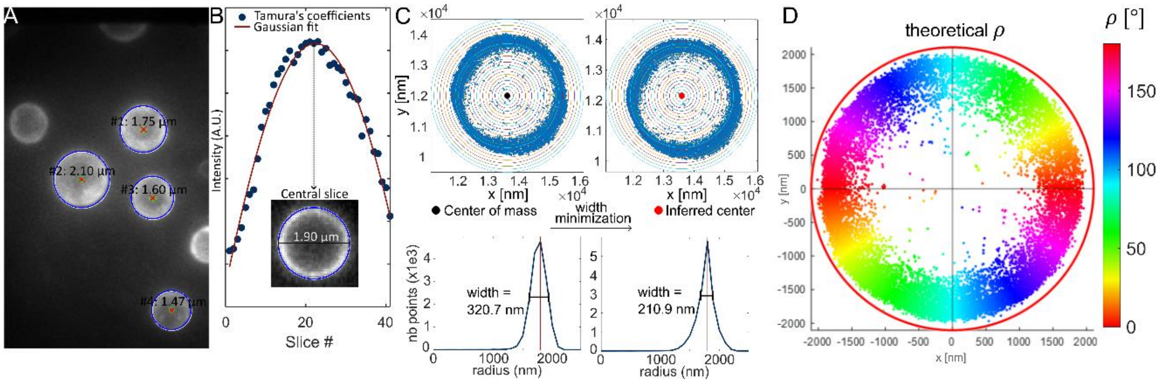


**Supplementary Figure S13. Outline of the analysis of SMOLM results of NR-labelled lipid bilayers covering silica beads.** A) The beads are identified on the maximum intensity projection image of a widefield z-stack (using Matlab function 'imfindcircles'). B) The central slice of the bead is inferred by computing Tamura's coefficients at each slice. The central slice position is given by the Gaussian fit of the Tamura's coefficients curve. The bead diameter is computed using the function 'imfindcircles' on the central slice of the z-stack. C) Top: The number of detections inside an annulus of increasing radius is computed. The width of the peak of this curve is minimized by varying the center of the annulus to find the center of the bead. D) Knowing the radius and the center of the bead, the theoretical orientation $\boldsymbol{\rho}$ (color-coded) can be computed for each localization.

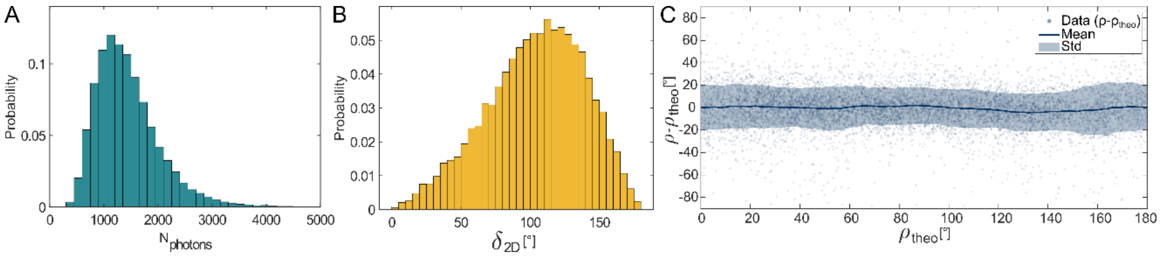


**Supplementary Figure S14. Quantifications on PolMFM imaging of NR labelling a SLB covering silica beads.** A) Histogram of number of photons for each detection. B) Histogram of retrieved wobbling $\boldsymbol{\delta_{2D}}$. C) Bias and error on orientation retrieval as a function of theoretical orientation $\boldsymbol{\rho_{theo}}$. In the scatter plots, each point represents a detection. The line is a rolling average over a sliding window of 15° sampled each 2°. Only molecules for which N_ph>1500 and $\boldsymbol{\delta_{2D} < 100}$° are kept. The theoretical orientation $\boldsymbol{\rho_{theo}}$ is inferred from the retrieved 2D localization and inferred bead center as specified in Supplementary Note 5.

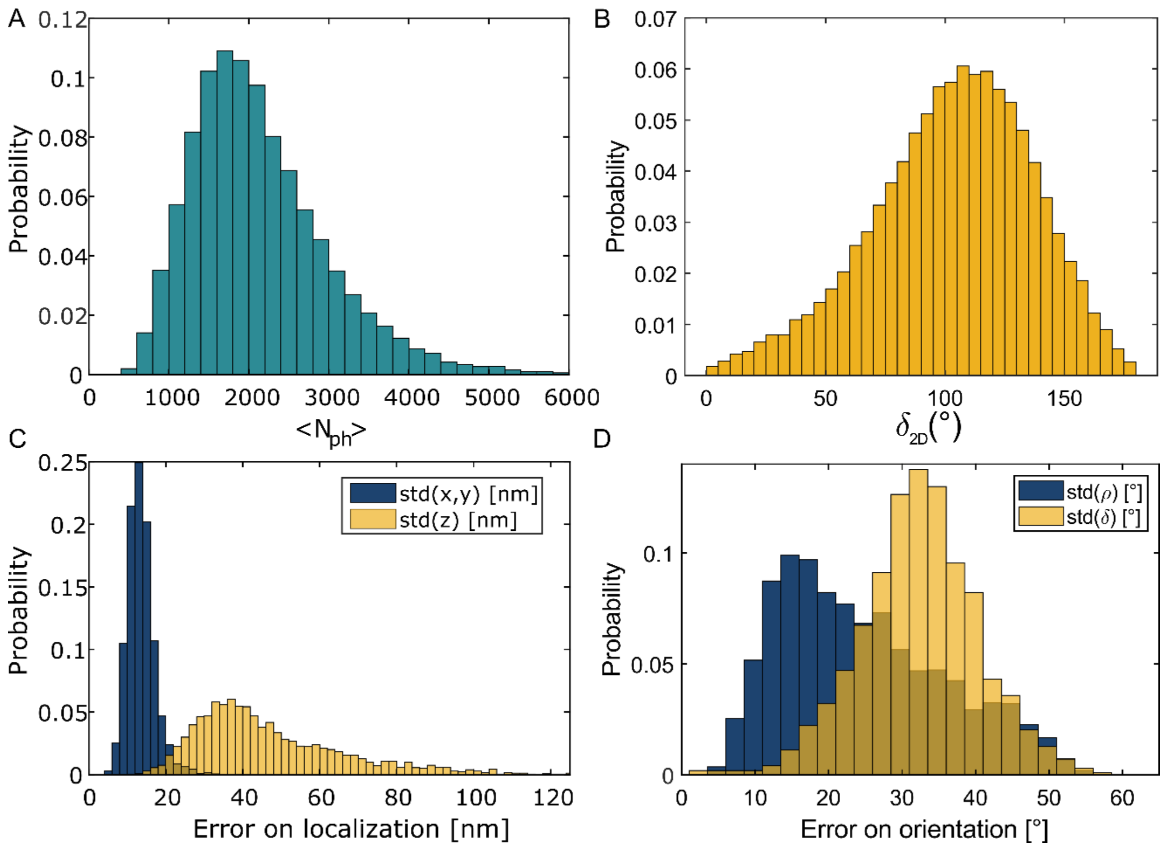


**Supplementary Figure S15. Quantifications on PolMFM imaging of AF568-phalloidin conjugates labelling actin in U2OS fixed cells.** A) Histogram of number of detected photons. B) Histogram of retrieved wobbling $\boldsymbol{\delta_{2D}}$. C) Histogram of error on lateral (blue) and axial (yellow) localization. The error is taken as the standard deviation of the retrieved value over multiple detections of the same molecule, where molecules appearing between 10 and 30 times are kept. D) Histogram of error on orientation $\boldsymbol{\rho}$ (blue) and projected wobbling $\boldsymbol{\delta_{2D}}$ (yellow). The error is taken as the standard deviation of the retrieved value over multiple detections of the same molecule, where molecules appearing at least 10 times are kept.

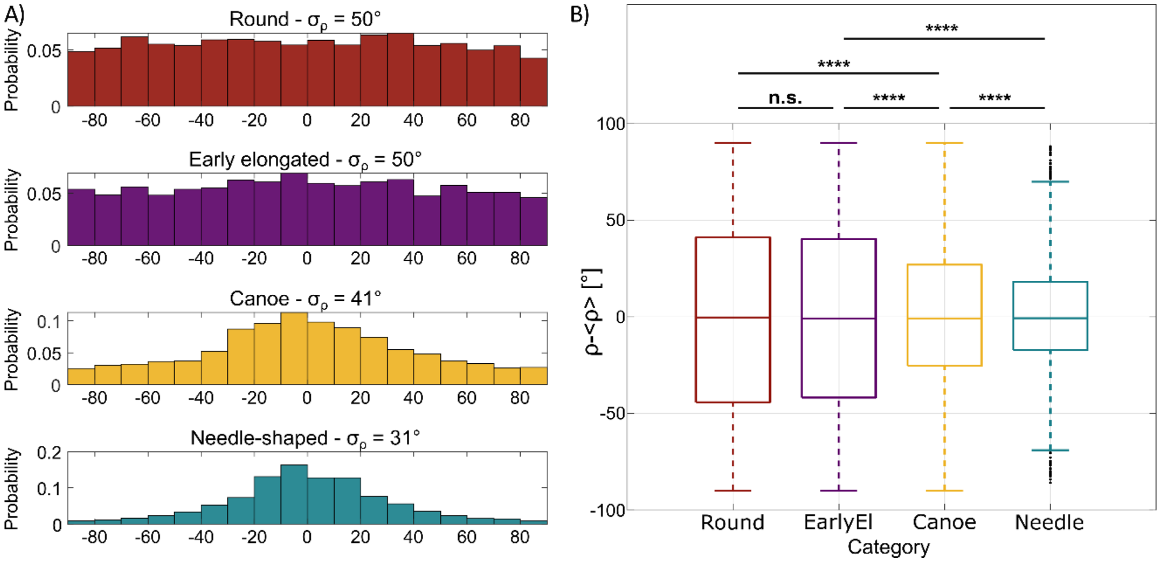


**Supplementary Figure S16. Distribution of orientations $\boldsymbol{\rho}$ at the different spermiogenesis stages.** A) Histograms of $\boldsymbol{\rho}$ distributions, centered to the mean value of each distribution, at 5 different identified stages of spermiogenesis. From top to bottom: round, early elongated, canoe, needle-shaped and mature. For each category, the distributions $\boldsymbol{\rho} - < \boldsymbol{\rho} >$ are concatenated from 4 to 7 different cells. B) Box plot of the distributions shown in A), with statistical tests (p-value from Levene's test) to compare the significance of variance change between the distributions. The number of molecules for each category is: 2809 (round), 2606 (early elongated), 1733 (canoe), 1037 (needle). No wobbling threshold was applied in this analysis.

**Supplementary Tables**

**Table S1.** Parameters used for PolMFM simulations.

| | |
|---|---|
| Refractive index sample | $n_2 = 1.33$ |
| Refractive index objective | $n_1 = 1.5$ |
| Numerical aperture | $NA = 1.4$ |
| Objective magnification | $OM = 100$ |
| Focal lens tube length | $f_{TL} = 200\ mm$ |
| Emission wavelength | $\lambda_{em} = 520\ nm$ |
| Interplane distance MFM | $\Delta z = 400\ nm$ |
| Central focal plane position | $d = \ -2.7\ \mu m$ |
| Relay lens focal length 1 | $f_1 = 150\ nm$ |
| Relay lens focal length 2 | $f_2 = 200\ nm$ |
| Camera pixel size | $px\ size_{ccd} = 16\ \mu m$ |
| Image pixel size | $px\ size = 120\ nm$ |

**Supplementary Information References**


1. Forkey, J. N., Quinlan, M. E. & Goldman, Y. E. Protein structural dynamics by single-molecule fluorescence polarization. *Progress in Biophysics and Molecular Biology* **74**, 1–35 (2000).
2. Backer, A. S. & Moerner, W. E. Determining the rotational mobility of a single molecule from a single image: a numerical study. *Opt. Express, OE* **23**, 4255–4276 (2015).
3. Lew, M. D., Backlund, M. P. & Moerner, W. E. Rotational Mobility of Single Molecules Affects Localization Accuracy in Super-Resolution Fluorescence Microscopy. *Nano Lett* **13**, 3967–3972 (2013).
4. Rimoli, C. V., Valades-Cruz, C. A., Curcio, V., Mavrakis, M. & Brasselet, S. 4polar-STORM polarized super-resolution imaging of actin filament organization in cells. *Nat Commun* **13**, 301 (2022).
5. Rimoli, C. V., Valades-Cruz, C. A., Curcio, V., Mavrakis, M. & Brasselet, S. 4polar-STORM polarized super-resolution imaging of actin filament organization in cells. *Nat Commun* **13**, 301 (2022).
6. Yan, T., Richardson, C. J., Zhang, M. & Gahlmann, A. Computational correction of spatially variant optical aberrations in 3D single-molecule localization microscopy. *Opt. Express, OE* **27**, 12582–12599 (2019).
7. Yan, T., Richardson, C. J., Zhang, M. & Gahlmann, A. Computational correction of spatially variant optical aberrations in 3D single-molecule localization microscopy. *Opt. Express, OE* **27**, 12582–12599 (2019).
8. Mertz, J. *Introduction to Optical Microscopy*. (Cambridge University Press, 2019).
9. Backer, A. S. & Moerner, W. E. Extending Single-Molecule Microscopy Using Optical Fourier Processing. *J. Phys. Chem. B* **118**, 8313–8329 (2014).
10. Hajj, B., Oudjedi, L., Fiche, J.-B., Dahan, M. & Nollmann, M. Highly efficient multicolor multifocus microscopy by optimal design of diffraction binary gratings. *Sci Rep* **7**, 5284 (2017).
11. Abrahamsson, S. *et al.* Fast multicolor 3D imaging using aberration-corrected multifocus microscopy. *Nat Methods* **10**, 60–63 (2013).

12. Hajj, B. *et al.* Whole-cell, multicolor superresolution imaging using volumetric multifocus microscopy. *Proc. Natl. Acad. Sci. U.S.A.* **111**, 17480–17485 (2014).
13. Thapa, V. vikas08thapa/Best_infocus_image_microscopy_stack. (2019).
